# Mitochondria-ER contacts in reactive astrocytes coordinate local perivascular domains to promote vascular remodelling

**DOI:** 10.1101/657999

**Authors:** Jana Goebel, Esther Engelhardt, Patric Pelzer, Vignesh Sakthivelu, Hannah M. Jahn, Milica Jevtic, Kat Folz-Donahue, Christian Kukat, Astrid Schauss, Christian K. Frese, Patrick Giavalisco, Alexander Ghanem, Karl-Klaus Conzelmann, Elisa Motori, Matteo Bergami

## Abstract

Astrocytes have emerged for playing important roles in brain tissue repair, however the underlying mechanisms remain poorly understood. We show that acute injury and blood-brain barrier disruption trigger the formation of a prominent mitochondrial-enriched compartment in astrocytic end-feet which enables vascular remodeling. Integrated imaging approaches revealed that this mitochondrial clustering is part of an adaptive response regulated by fusion dynamics. Astrocyte-specific conditional deletion of Mitofusin 2 (*Mfn2*) suppressed perivascular mitochondrial clustering and disrupted mitochondria-ER contact sites. Functionally, two-photon imaging experiments showed that these structural changes were mirrored by impaired mitochondrial Ca^2+^ uptake leading to abnormal cytosolic transients within end-feet *in vivo*. At the tissue level, a compromised vascular complexity in the lesioned area was restored by boosting mitochondrial-ER perivascular tethering in MFN2-deficient astrocytes. These data unmask a crucial role for mitochondrial dynamics in coordinating astrocytic local domains and have important implications for repairing the injured brain.

## Introduction

Astrocytes regulate essential aspects of brain energy metabolism (Belanger et al., 2011) but also play important roles in the progression and possible resolution of numerous brain pathologies, including traumatic brain injury and stroke (Sofroniew, 2015). These types of injury often result in significant damage to the cerebrovasculature and are usually accompanied by blood-brain barrier breakdown, intracerebral hemorrhage, hypoxia, secondary inflammation and neurodegeneration (Prakash and Carmichael, 2015; Salehi et al., 2017). While a number of factors concerning the severity of the primary insult contribute to the extent of tissue damage and thus influence the subsequent attempt to repair, our understanding of the mechanisms underlying neovascularization in the injured area and which exact cellular components are recruited is still rudimentary.

Besides endothelial cells and pericytes, which constitute the actual blood-brain-barrier, astrocytic end-feet functionally ensheathe most of the cerebrovascular network and serve as specialized dynamic exchange sites for ions, water and energy substrates with brain parenchyma (Iadecola, 2017). While maintenance of this tight coupling ensures the supply of metabolites across the gliovascular interface, thereby contributing to neurovascular coupling (Iadecola, 2017), the structural and functional changes experienced by astrocytic perivascular end-feet following injury and the ensuing vascular damage are much less understood. In these settings, astrocytes are known to acquire a reactivity cellular state which may underlie both beneficial and deleterious functions (Khakh and Sofroniew, 2015; Liddelow and Barres, 2017). Interestingly, some of these functions have been described for regulating angiogenesis via the secretion of trophic factors and molecules which can ultimately lead to vascular remodeling (Salehi et al., 2017). Furthermore, evidence for a prominent physical association between perivascular astrocytes and vessels in the peri-lesioned area has been reported following acute injury, particularly at a time matching with the formation of new vessels (Horng et al., 2017; Villapol et al., 2014), thus suggesting that structural changes at the gliovascular interface may be critical in regulating vascular remodeling after injury. Importantly, while emerging evidence indicates that the contribution of astrocytes to disease progression depends on the specific type of reactivity state acquired (Anderson et al., 2016; Liddelow et al., 2017), it is becoming clear that these states underlie not only major changes in morphology and gene expression but also a significant extent of metabolic plasticity (Chao et al., 2019; Polyzos et al., 2019). Supporting this notion, astrocytes can utilize oxidative phosphorylation (OXPHOS) for their energy metabolism (Ignatenko et al., 2018; Lovatt et al., 2007), yet they efficiently sustain for long periods of time glycolytic fluxes (Supplie et al., 2017), underscoring the capability of these cells to accommodate a significant metabolic rewiring depending on substrate availability and local energy needs (Hertz et al., 2007). This important form of plasticity is emphasized by the fact that astrocytes reacting to injury *in vivo* can adjust their metabolic signature by efficiently and reversibly modifying the architecture of their mitochondrial network (Motori et al., 2013; Owens et al., 2015), i.e. the central hub for cellular energy metabolism and metabolic signaling, thus suggesting that these mitochondrial responses may also represents an important mechanism whereby astrocytes actively contribute to tissue remodeling.

The architecture of the mitochondrial network in cells is usually very dynamic and its maintenance depends upon regulated fusion-fission events as well as on abundant contact sites with the ER and other organelles (Labbe et al., 2014). In mammalian cells, the main drivers of mitochondrial membrane dynamics are mitofusins (MFN1 and MFN2) (Chen et al., 2003) and optic atrophy-1 (OPA1) (Cipolat et al., 2004) for mitochondrial fusion, while dynamin-related protein-1 (DRP1) is the key player in outer mitochondrial fission (Ishihara et al., 2009). Together, the coordinated action of these molecules shapes mitochondria towards more fragmented or elongated morphologies to match precise cellular metabolic needs (Dietrich et al., 2013; Gomes et al., 2011; Rambold et al., 2011). Functionally, this mitochondrial remodeling is also regulated by a physical tethering with ER membranes to form specialized contact sites (so-called mitochondria-associated membranes or MAMs) that control important metabolic signaling functions (Scorrano et al., 2019), including lipid trafficking as well as the formation of Ca^2+^ and ROS microdomains (Csordas et al., 2018). Intriguingly, evidence exists for complex mitochondrial and ER morphologies in astrocytes *in situ*, where these organelles have been found to reach fine perisynaptic processes and end-feet (Gobel et al., 2018; Jackson and Robinson, 2018; Lovatt et al., 2007; Mathiisen et al., 2010; Motori et al., 2013). While this spatial distribution suggests the direct contribution of MAMs to specific astrocytic functions, whether and to which extent a dynamic remodeling of these two organelles may effectively couple the acquisition of a reactive state with functional metabolic changes is unclear.

Here, we provide evidence that acute brain injury triggers a distinctive clustering of mitochondria in perivascular astrocytic end-feet, where they form extensive contact sites with the ER. Our data indicate that this clustering is coordinated by mitochondrial fusion dynamics and generates a local mitochondrial-enriched domain surrounding microvessels. *Mfn2* deficiency in reactive astrocytes prevented injury-induced perivascular accumulation of mitochondria, altered the extent of mitochondria-ER tethering leading to disrupted Ca^2+^ dynamics in astrocyte end-feet, and ultimately impaired angiogenesis and vascular remodeling in the injured area. Importantly, our data indicate that vascular remodeling can be restored in absence of mitochondrial fusion by forcefully enhancing perivascular mitochondria-ER contact sites. These results establish a mechanism for mitochondrial fusion in orchestrating local functional domains in astrocytes *in vivo* and unravel a key role for astrocytic mitochondria-ER contact sites in sustaining microvasculature remodeling during repair.

## Results

### Astrocyte end-feet are naturally enriched in mitochondria-ER contact sites

In order to investigate how the architecture of mitochondrial and ER networks may match the morphological complexity of astrocytes we utilized a virus-based strategy to label specifically these organelles *in vivo* (Figure 1A). Minimal amounts of either hGFAP promotor-driven adeno-associated viruses (AAV) or modified EnvA-pseudotyped rabies viruses (RABV) were stereotactically injected into the cortex of wild-type or hGFAP-TVA mice, respectively, to drive the expression of mitochondrial- or ER-targeted fluorophores (i.e. mitoRFP and ER-GFP). Both viral-based approaches have been previously shown to efficiently restrict the expression of transgenes to astrocytes in the adult brain (Motori et al., 2013; Shigetomi et al., 2013). Single-astrocyte analysis one week after virus delivery revealed a complex morphology of these organelles, which were found decorating the most peripheral astrocytic processes, including fine branchlets (Figure 1B). Interestingly, besides few primary branches originating from the soma, structures identified as perivascular end-feet (i.e., possessing a tube-like morphology and surrounding CD31+ vessels) were often enriched in ER and mitochondria (Figure 1C and S1A-B). In these regions, the ER appeared to virtually delineate the shape of vessels, while mitochondria often formed a dense meshwork that was much similar to the one observed within primary branches rather than distally-located fine perisynaptic processes (branchlets) (Figure 1C and S1A-B). Experiments conducted by labelling the microvasculature via systemic dextran injection prior to sacrifice revealed that astrocytic ER and mitochondrial networks outlined the labelled vessels to the extent that often whole sections of the microvasculature appeared wrapped by a thin but discernible layer of astrocytic organelles (Figure 1D-E). In contrast, virus-mediated labelling of other organelles including peroxisomes and lysosomes yielded a much different distribution, being largely confined to the cell body and major branches (Figure S1C-F).

**Figure 1.**
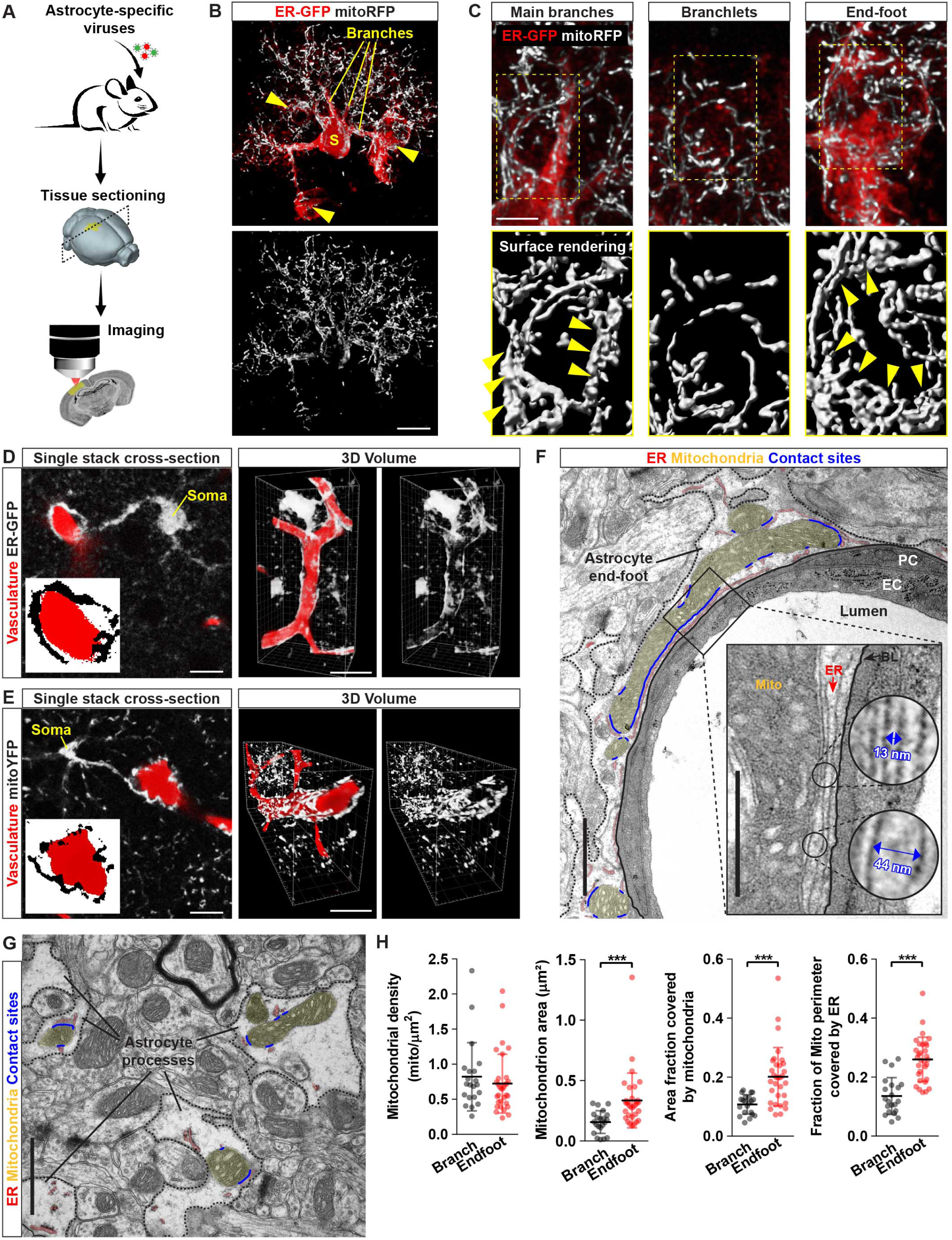
Astrocytic end-feet are enriched in mitochondria-ER contact sites. **(A)** Experimental design used to express organelle-targeted fluorescent sensors in astrocytes *in vivo*. **(B)** Example of a cortical astrocyte co-transduced with ER-GFP and mitoRFP viruses. Yellow arrowheads point to the end-feet. Bar, 10 µm. **(C)** Magnifications of the astrocyte shown in B. Yellow arrowheads point to bundles of elongated mitochondria. Bar, 5 µm. **(D-E)** Examples of astrocytes transduced with ER-GFP (D) or mitoYFP (E) wrapping around dextran-labeled vessels. Insets show zooms of the perivascular end-foot. Side panels show a 3D rendering of the same astrocytes. Bars, 10 and 25 µm. **(F)** EM picture of a vessel cross-section showing the astrocytic end-foot (segmented black line) and its organelles (mitochondria: yellow; ER: red; contact sites: blue). The inset shows mitochondria-ER contact sites lining the basal lamina. Bars, 2 and 1 µm. **(G)** EM picture of perisynaptic astrocytic processes and their organelles. Bar, 2 µm. **(H)** Quantification of mitochondrial parameters in branches (n= 21 vessel cross-sections from 3 mice) and end-feet (n= 32 vessel cross-sections from 3 mice; nonparametric Mann-Whitney t-test). ***, p< 0.001. PC: pericyte; EC: endothelial cell; BL: basal lamina. See also Figure S1.

At the ultrastructural level, astrocytic end-feet appeared enriched with ER membranes surrounding not only the basal lamina but also most of mitochondria located within the perivascular process (Figure 1F and Figure S1G-H). In particular, substantial portions of the mitochondrial perimeter were bordered by ER membranes and, at these contact sites, the two organelles maintained an average reciprocal distance of 18.9 ± 5.0 nm (Figure 1F). By comparison, both the size of mitochondria and the extent of ER membranes were smaller in perisynaptic astrocytic processes, resulting in visibly fewer contact sites, despite a similar mitochondria-ER average distance of 20.4 ± 7.0 nm (Figure 1G). Accordingly, morphological quantification revealed a net enrichment in mitochondrial area and mitochondria-ER tethering domains within the end-feet (Figure 1H), suggestive of key metabolic functions being regulated by these two organelles at perivascular sites.

### Marked remodelling of astrocyte mitochondrial networks following cortical injury

Astrocyte reactivity states are characterized by prominent changes in energy metabolism and mitochondrial network morphology (Castejon, 2015; Hamby et al., 2012; Motori et al., 2013; Zamanian et al., 2012), raising the question of whether perivascular organelle distribution may become affected during the acquisition of a reactive cellular state. To answer this question, we utilized a genetic approach to conditionally express mitoYFP in adult astrocytes and investigate in detail mitochondrial morphology after cortical stab-wound (SW)-injury *in vivo* (Figure 2A). Human-GFAP-CreER mice (Chow et al., 2008) were crossed with mitoYFP floxed-stop mice (Sterky et al., 2011) and the resulting line was induced with tamoxifen at the age of 6-8 weeks. With this approach, about 88% of cortical astrocytes (S100β+) underwent recombination (Figure S2A), allowing for a systematic analysis of the changes in the mitochondrial network of cells located in the vicinity of the lesion track (i.e. the area mostly enriched in extravasating pro-inflammatory CD45+ leukocytes) (Figure 2B and S2B). In particular, by one week following SW, astrocytes reacted by overt mitochondrial fragmentation throughout all their cellular territories (Figure 2C and S2B) despite no major changes in the overall expression levels of mitochondrial fission-fusion proteins detected at this time (Figure S2F-G), suggesting the occurrence of post-translational modifications of the existing fission-fusion protein machinery (Anton et al., 2013; Motori et al., 2013). Yet, detailed morphometric analysis of reconstructed whole mitoYFP+ astrocytes revealed that, irrespective of their “metabolic” state (i.e., whether resting or reactive), the mitochondrial network in these cells was usually composed by highly heterogeneous morphologies, with both tubular and very long (up 8-10 μm) as well as much shorter organelles (less than 0.5 μm) (Figure 2C). This morphological diversity became apparent when plotting the length versus sphericity of the whole mitochondrial population of several reconstructed astrocytes selected for their close proximity to the lesion track (Figure 2D): by 7 days post-SW the mitochondrial network displayed a significant shift towards fragmentation with over 60% of the whole mitochondrial population being <1 μm in length, in contrast to a 43.5% in control astrocytes (Figure 2D). Whole-cell, time-course analysis during a period ranging from 3 days to 2 months after SW-injury revealed that while the fraction of fragmented mitochondria sharply increased during the first week, the network was restored to levels comparable to control astrocytes by the third week (Figure 2E). This trend was mirrored by opposite changes in the proportion of tubular mitochondria, confirming that the evolving reactive state of astrocytes proximal to the lesion is accompanied by a time-dependent remodelling of the whole mitochondrial network over the course of several weeks after injury (Motori et al., 2013).

**Figure 2.**
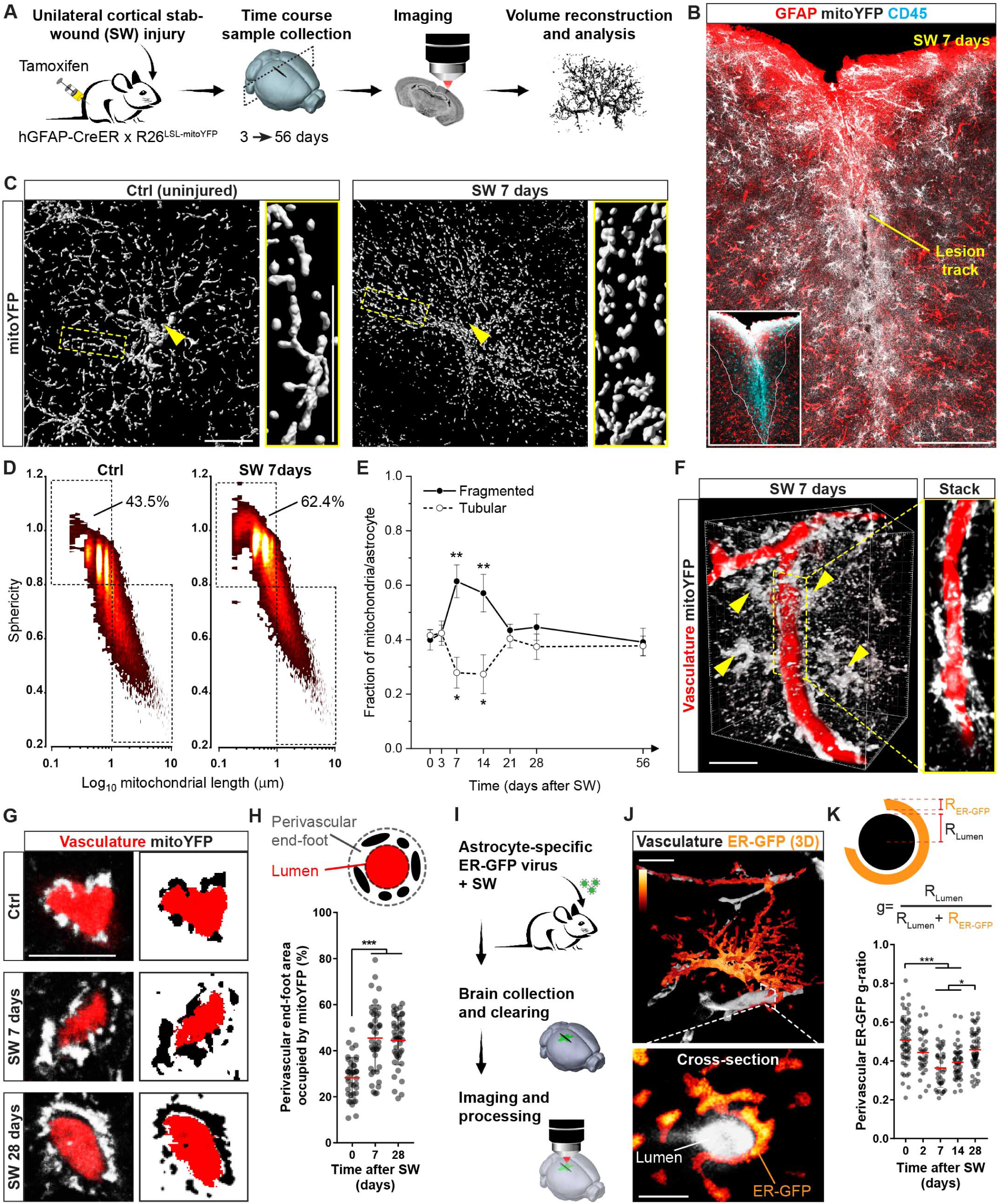
Dynamic remodelling of astrocyte mitochondrial and ER networks following injury. **(A)** Experimental design for examining the mitochondrial network in astrocytes *in vivo*. **(B)** Example of an hGFAP::CreER x R26^LSL-mitoYFP^ mouse at 7 days after cortical SW-injury. The inset shows extravasating CD45+ leukocytes in the lesion core. Bar, 150 µm. **(C)** Surface rendering of mitochondrial networks in control astrocytes (uninjured animals) or in reactive astrocytes proximal to the lesion track. Yellow arrowheads point to the soma. Zooms depict the predominant network morphology in peripheral branches. Bar, 15 µm. **(D)** Density plots depicting the morphological heterogeneity of the mitochondrial population in individual astrocytes under resting (Ctrl, uninjured animals) or reactive conditions (SW 7days). The proportion of fragmented mitochondria based on threshold values for mitochondrial sphericity (0.8) and length (1 µm) is shown. **(E)** Time-course analysis of mitochondrial fragmentation quantified as in D (n≥ 3 mice/time point, with 8-15 astrocytes/mouse; one-way ANOVA followed by Dunnett’s post-hoc test). **(F)** Volume reconstruction of mitoYFP+ astrocytes (arrowheads) surrounding dextran-labelled vessels at 7 days post-SW. A single-stack is shown. Bar, 25 µm. **(G)** Examples of vessel cross-sections showing perivascular astrocytic mitoYFP in control (uninjured animals) and injured conditions. Bar, 10 µm. **(H)** Quantification of perivascular mitoYFP density displayed as area fraction (n≥ 30 vessels/time-point; nonparametric Kruskal-Wallis test). **(I)** Experimental design for analyzing the astrocytic ER. **(J)** 3D example of an astrocyte expressing ER-GFP (signal density shown in pseudocolors). Bars, 10 and 5 µm. **(K)** Quantification of the ER-GFP perivascular *g-ratio* at the indicated time-points (n≥ 35 vessels/time-point; nonparametric Kruskal-Wallis test). **, p < 0.01, ***, p < 0.001. See also Figure S2.

Interestingly, inspection of microvessels proximal to the lesion (labelled via either dextran injection or CD31 immunostaining) revealed a conspicuous accumulation of astrocytic mitochondria in perivascular end-feet (Figure 2F and S2C). In particular, analysis of vessel cross-sections disclosed that the extent of mitochondria surrounding the vessels markedly increased by 7 and 28 days after SW (Figure 2G-H), the latter being a time when mitochondrial network morphology astrocyte-wide had already normalized back to control levels (Figure 2E). In contrast, mitochondrial density in peripheral branches and total mitochondrial mass in astrocytes (the latter examined both via microscopic mitoYFP quantification and label-free proteomic analysis of markers associated with mitochondrial biogenesis and mass in sorted astrocytes) appeared only mildly affected (Figure S2D-E and S2H). We next assessed whether the ER may also undergo a similar extent of remodelling in response to injury. Reactive astrocytes expressing ER-GFP appeared to retain a significant amount of ER at perivascular end-feet surrounding CD31+ vessels (Figure S2C). Three-dimensional reconstruction of individual ER-GFP-expressing astrocytes in conjunction with dextran-labelling revealed the whole distribution of the ER network across distinct astrocytic territories in uninjured hemispheres (Figure 2J and Figure S2I). In these control samples, the GFP signal allowed for the assessment of a perivascular ER-GFP “*g*-ratio” to investigate changes in perivascular ER dynamics and normalize these to putative variations in microvessel diameter (Figure 2K and S2J). This analysis disclosed a time-dependent increase in the thickness of perivascular ER-GFP signal, which peaked by 7 days post-SW but reverted to near-basal conditions by 28 days (Figure 2K). These results were corroborated relative volume distribution analysis of the ER-GFP signal (i.e. signal density) across astrocytic compartments (Figure S2I). In control astrocytes, perivascular end-feet accounted for 19.2% of all ER-GFP signal in individual cells (Figure S2L). In contrast, in injury-induced reactive astrocytes an accumulation of ER-GFP signal was observed in the end-feet (35.9%) at the expenses of main branches (where the relative ER-GFP proportion decreased from 39.1% in controls to 25.2% in injured samples) (Figure S2K and S2L-M). Interestingly, by 28 days after SW the relative distribution of ER-GFP signal mostly normalized (Figure S2K and S2L-M), suggesting that in contrast to the enduring response of the mitochondrial network in perivascular end-feet (Figure 2H), remodelling of the ER compartment may only be temporary. Together, these data reveal that mitochondrial and ER networks undergo a regionalized morphological rearrangement in perivascular end-feet of astrocytes reacting to acute injury.

### Conditional deletion of *Mfn2* disrupts perivascular mitochondria-ER contact sites in astrocytes

The reversible transition of the mitochondrial network from fragmentation at 7 days post-SW to a tubular network by 21 days (Figure 2E), together with the peculiar remodelling of mitochondria within the end-feet of reactive astrocytes (Figure 2G-H), argues in favour of regulatory mechanisms playing a role in adjusting the mitochondrial network to match evolving metabolic needs in response to injury. We reasoned that interfering with these mechanisms by preventing mitochondrial re-tubulation may provide a valid approach to dissect the specific role of this network remodelling for astrocyte physiology (Figure 3A). We opted for the conditional deletion of the GTPase protein MFN2, which is a key effector of mitochondrial outer membrane fusion dynamics but also plays a role in maintaining mitochondria-ER tethering domains (de Brito and Scorrano, 2008). Specific deletion in astrocytes was achieved by crossing Mfn2 floxed mice (Lee et al., 2012) with the inducible hGFAP-CreER x mitoYFP floxed-stop mouse line (hereafter defined as Mfn2^cKO^ mice). Few weeks after tamoxifen-mediated recombination induced in 2-month old mice, *Mfn2* gene knock-out was assessed by genotyping of isolated brain cortices (Figure S3A-B) and protein depletion was validated via mass spectrometry analysis of astrocytes acutely sorted from brain cortex via magnetic cell separation (MACS) (Figure 3B). In contrast to classic astrocytic markers (i.e., GLAST, GLT-1, ALDH1L1 and AQP4) or other reference mitochondrial proteins (OPA1 and TOMM40), MFN2 was specifically and markedly downregulated (more than 9 folds, Figure 3B). Analysis of transmission electron microscopic (TEM) pictures revealed fewer and circular mitochondria of significant size within the end-feet of Mfn2^cKO^ astrocytes, in net contrast to Mfn2^WT^ samples, in which elongated and branched morphologies were observed lining the basal lamina of microvasculature cross-sections (i.e., having an average vessel diameter of 3.5 ± 0.6 μm in Mfn2^WT^ and 3.6 ± 0.6 μm in Mfn2^cKO^) (Figure 3C and 3E). Close inspection of perivascular end-feet revealed however that the overall distribution of the ER was not overtly affected in Mfn2^cKO^ astrocytes, with long stretches of ER tubule surrounding the basal lamina as in Mfn2^WT^ astrocytes (Figure 3D). Interestingly, Mfn2^cKO^ mitochondria were less enriched in ER contact sites despite the nearby presence of abundant ER membranes (Figure 3C-E). Notably, deletion of *Mfn2* in astrocytes did not visibly affect mitochondrial cristae morphology within the examined time frame (4 weeks post-tamoxifen treatment) (Figure 3C). Together, these results indicate that conditional deletion of *Mfn2* in adult astrocytes *in vivo* leads to ultrastructural morphological changes of their mitochondria and a concomitant reduction in the extent of mitochondria-ER contact sites within end-feet.

**Figure 3.**
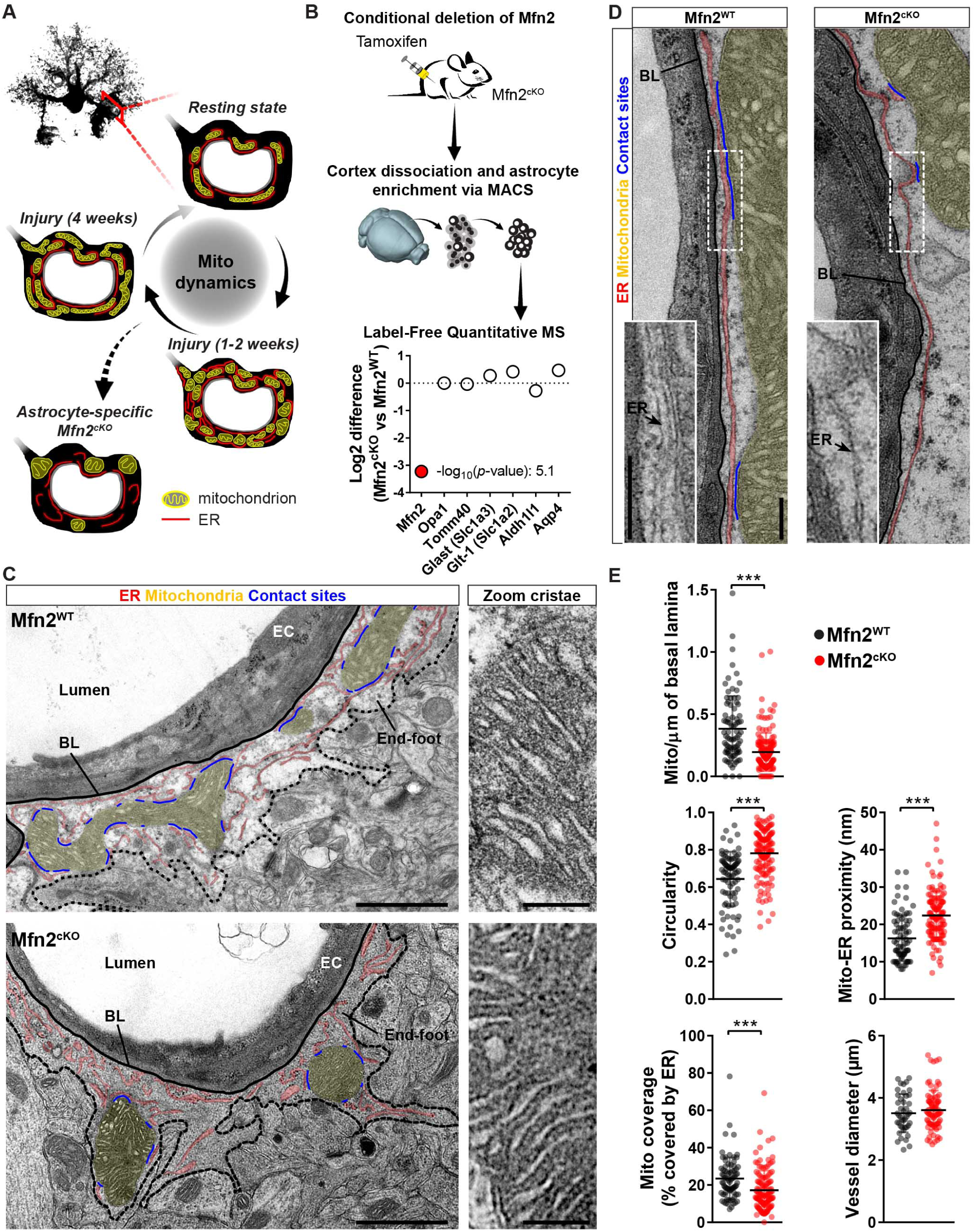
*Mfn2* deletion affects astrocytic mitochondria-ER tethering domains. **(A)** Proposed model showing the extent of mitochondrial remodeling taking place in astrocytic end-feet during injury and the expected phenotype following *Mfn2* deletion. **(B)** Experimental design for validating *Mfn2* knock-out in astrocytes by MACS enrichment and proteomic analysis. The plot shows MFN2 protein abundance in Mfn2^cKO^ samples compared to other mitochondrial and classic astrocytic markers (n= 4 Mfn2^cKO^ mice and 3 Mfn2^WT^ mice). **(C)** EM pictures of astrocytic end-feet in Mfn2^WT^ and Mfn2^cKO^ mice at 4 weeks post-tamoxifen treatment. Mitochondria and ER contact sites are highlighted in different colors. Right panels depict zooms of mitochondrial cristae. EC: endothelial cell; BL: basal lamina. Bars, 1 µm and 200 nm. **(D)** Details of astrocytic end-feet showing the perivascular distribution of ER tubules and their contact sites with mitochondria in Mfn2^WT^ and Mfn2^cKO^ mice. Bars, 250 nm. **(E)** Quantification of the indicated ultrastructural parameters in Mfn2^WT^ (n= 85 vessel cross-sections from 4 mice) and Mfn2^cKO^ perivascular end-feet (n= 145 vessel cross-sections from 3 mice; non-parametric Mann-Whitney t-test). ***, p < 0.001. See also Figure S3.

### Astrocyte-specific *Mfn2* deletion abrogates perivascular remodelling of still functional mitochondria

We next asked the question whether *Mfn2* deletion would be sufficient to prevent astrocyte mitochondrial network remodelling in response to acute injury. Histological and protein examination of astrocytes (i.e., via label-free proteomic analysis of sorted reactive astrocytes at 4 weeks post-SW) derived from lesioned Mfn2^cKO^ animals revealed no overt abnormalities in the extent of GFAP or Vimentin expression (i.e., classic markers of reactivity) within the area surrounding the lesion track at 7 days post-SW (Figure 4A and S3C-D). Analysis of recently annotated additional markers of astrocytic reactivity (Liddelow et al., 2017) detected in our proteomic dataset revealed variable changes in their expression levels, with no obvious trend towards a higher or lower reactivity state (Figure S3D). At the single-cell level, however, mitochondrial network morphology in Mfn2^cKO^ astrocytes appeared significantly affected even in uninjured conditions when compared to control astrocytes (Figure S4A). In particular, mitochondria appeared fragmented throughout astrocytic territories, confirming loss of MFN2 and the consequent lack of mitochondrial fusion dynamics starting as soon as one week after tamoxifen-induced recombination. In contrast, the ER network retained an overall intact morphology in the absence of MFN2 (Figure S4B). Interestingly, conditional deletion of *Mfn1* resulted in somewhat heterogeneous and less pronounced morphological changes (Figure S4A), suggesting either differences in the relative expression levels of the two mitofusins or potential compensatory effects in the expression levels of MFN2 following *Mfn1* deletion (Figure S3E), as previously reported for other tissues (Kulkarni et al., 2016). Single-cell, time-course analysis of mitochondrial morphology revealed that both Mfn2^cKO^ and Mfn1^cKO^ astrocytes retained the capability to undergo further fragmentation following SW (Figure 4B). In particular, by 7 days post-SW, i.e. at the peak of fragmentation in control astrocytes, the overall proportion of fragmented versus tubular mitochondria appeared almost indistinguishable between examined groups (Figure 4B). However, while control and Mfn1^cKO^ astrocytes gradually and efficiently reformed a tubular network by 28 days post-SW, Mfn2^cKO^ astrocytes lacked this ability and were left with visibly fragmented mitochondria (Figure 4B-C). Importantly, perivascular mitochondrial clustering induced by injury was significantly impaired in Mfn2^cKO^ astrocytes proximal to the lesion site, in contrast to wild-type (control) and Mfn1^cKO^ astrocytes, in which the extent of mitoYFP signal essentially doubled (Figure 4D-E and S4C). Conspicuously, TEM analysis of Mfn2^cKO^ astrocytes confirmed a marked reduction in mitochondrial density and mitochondria-ER contact sites in perivascular end-feet despite intact mitochondrial cristae and presence of abundant ER tubules (Figure S4D-F).

**Figure 4.**
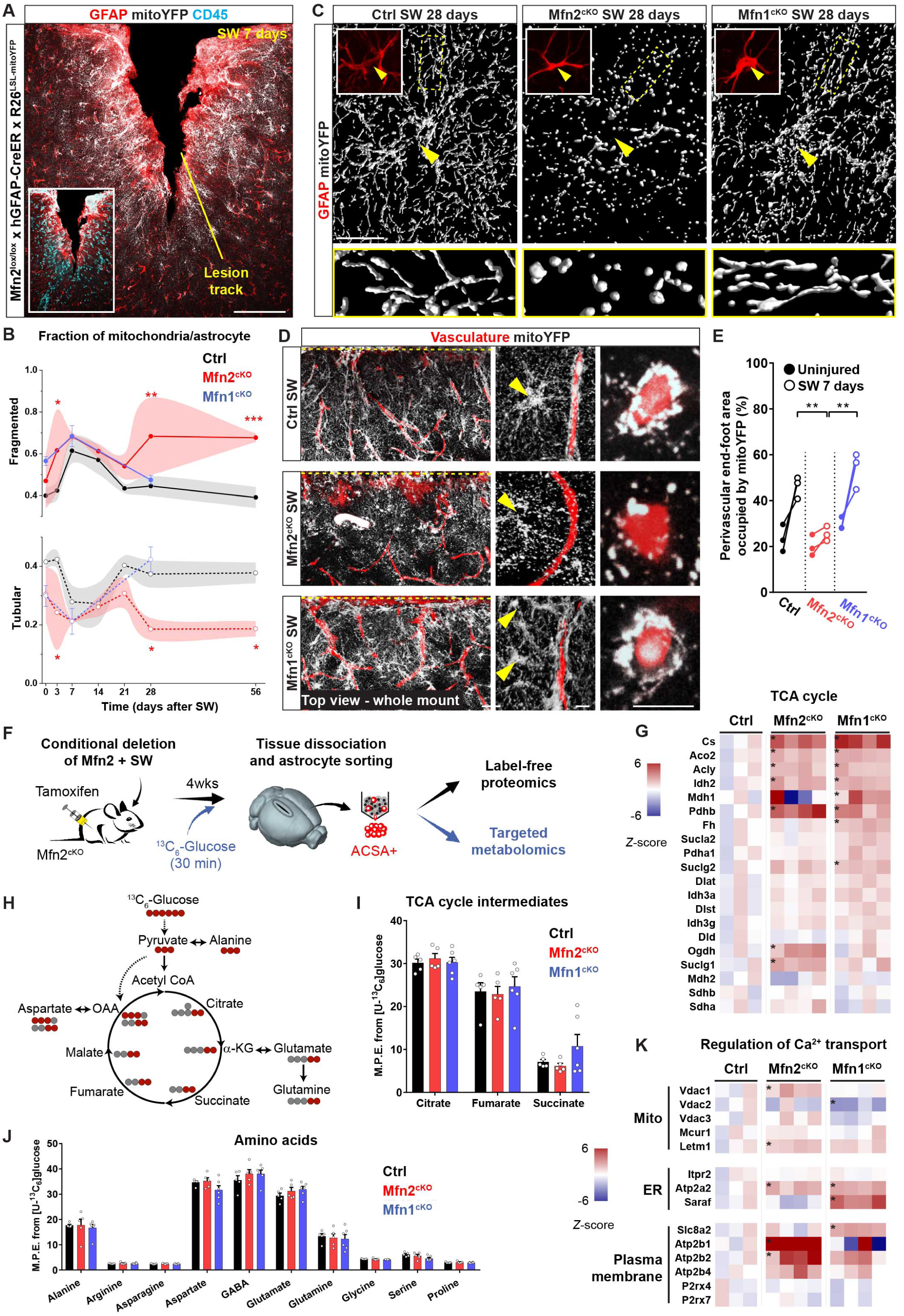
*Mfn2* deletion in adult reactive astrocytes prevents perivascular clustering of still functional mitochondria. **(A)** Example of cortical SW-injury in Mfn2^cKO^ mice at 7 days. The inset shows CD45+ leukocytes within the lesion core. Bar, 100 µm. **(B)** Time-course analysis of mitochondrial fragmentation in Mfn2^cKO^ and Mfn1^cKO^ astrocytes (n≥ 3 mice/time point, 8-15 astrocytes/mouse; two-way ANOVA followed by Tukey’s post-hoc test). **(C)** Examples of mitochondrial morphologies in astrocytes (arrowheads point to soma) proximal to the lesion site at 28 days post-SW. Zooms on the right depict peripheral branches. Insets show immunoreactivity for GFAP. Bar, 20 µm. **(D)** Top view projections (100 μm deep) of Mfn2^cKO^ and Mfn1^cKO^ whole-mount injured cortices (7 days) following tissue clearing. The penetrating SW-injury site (perpendicular to the view) is indicated by a yellow dashed line on top of each panel. Middle panels depict a mitoYFP-expressing astrocyte proximal to the lesion track and nearby vessels. Right panels depict a vessel cross-section. Bars, 50, 10 and 10 µm. **(E)** Quantification of astrocytic mitoYFP perivascular density (n= 3 mice/condition, with a total of at least 80 vessel sections quantified; the contralateral uninjured sides were utilized as internal controls; one-way ANOVA followed by Holm-Sidak’s post-hoc test). **(F)** Schematic illustrating the experimental protocol used for astrocyte isolation via anti-ACSA staining and cell sorting followed either by proteomic analysis. A similar approach was used to perform targeted metabolomics of mice supplied with ^13^C_6_-Glucose. **(G)** Heat maps of normalized LFQ intensities of TCA cycle and associated enzymes in reactive astrocytes of Mfn2^cKO^ and Mfn1^cKO^ mice. Values are color-coded according to their z-score. Significant protein changes (-log_10_ of the *p*-value ≥1.3) are indicated with an asterisk (n= 4 Mfn2^cKO^ mice, 4 Mfn1^cKO^ mice and 3 Ctrl mice). **(H)** Atom-resolved map of the expected main isotope distribution after ^13^C labeling (red dots indicate ^13^C atoms) in intermediates of the TCA cycle following supplementation of ^13^C_6_-Glucose (delivered by systemic injection into mice 30 minutes before sacrifice). **(I)** Relative enrichment (M.P.E) in ^13^C-labeled species for each of the indicated TCA cycle intermediates at 4 weeks after SW (n= 5 Mfn2^cKO^ mice, 6 Mfn1^cKO^ mice and 5 Ctrl mice; two-way ANOVA followed by Dunnett’s test). **(J)** Relative enrichment in ^13^C-labeled species for each of the indicated amino acids at 4 weeks after SW (n= 5 Mfn2^cKO^ mice, 6 Mfn1^cKO^ mice and 5 Ctrl mice; two-way ANOVA followed by Dunnett’s test). **(K)** Heat maps of normalized LFQ intensities of proteins regulating Ca2+ transport across the indicated organelles in reactive astrocytes of Mfn2^cKO^ and Mfn1^cKO^ mice. Values are color-coded according to their z-score. Significant protein changes (-log_10_ of the *p*-value ≥1.3) are indicated with an asterisk (n= 4 Mfn2^cKO^ mice, 4 Mfn1^cKO^ mice and 3 Ctrl mice). *, p < 0.05, **, p < 0.01, ***, p < 0.001. See also Figure S3 and S4.

The presence of intact cristae structure in reactive Mfn2^cKO^ astrocytes raised the question of whether these mitochondria were still metabolically competent. We thus further examined our proteomic dataset of acutely sorted Mfn2^cKO^ astrocytes at 4 weeks post-SW (Figure 4F and S3C). Ingenuity Pathway Analysis (IPA) of our dataset disclosed the Oxidative Phosphorylation pathway among the Mfn2^cKO^-specific, down-regulated hits in our samples (Figure S3F), yet detailed inspection of mitochondrial respiratory chain complexes indicated that only a few of the detected subunits in complexes I, III, IV and V were significantly down-regulated (Figure S3G). Likewise, proteins associated with mitochondrial stress responses revealed that only few of them were significantly up-regulated in Mfn2^cKO^ astrocytes (Figure S3H), suggesting that absence of MFN2 brings about only a modest mitochondrial dysfunction on top of potential changes induced by injury itself. Interestingly, we observed a general up-regulation in the protein expression levels of enzymes associated to the tricarboxylic acid (TCA) cycle (Figure 4G) and the catabolism of amino acids and their derivatives (Figure S3F), which have emerged as hallmarks of mitochondrial metabolic rewiring in multiple cell types (Chen et al., 2018). Of note, a similar upregulation was found in Mfn1^cKO^ astrocytes (Figure 4G and S3F). However, targeted metabolomics of sorted astrocytes following systemic infusion of ^13^C_6_-Glucose (Figure 4F) revealed no changes in the incorporation of glucose-derived carbon into TCA cycle intermediates or amino acids between control, Mfn2^cKO^ and Mfn1^cKO^ astrocytes (Figure 4H-J), indicating that mitochondrial bioenergetics are not overtly compromised in reactive Mfn2^cKO^ astrocytes up to 4 weeks post-SW.

Altogether, these results indicate that while conditional *Mfn2* deletion in reactive astrocytes prevents perivascular enrichment of mitochondria and mitochondria-ER contact sites, mitochondrial cristae structure and function remain to large degree unaffected after injury.

### Lack of MFN2 dampens astrocytic mitochondrial Ca^2+^ uptake and leads to abnormal perivascular Ca^2+^ transients after SW-injury *in vivo*

The absence of a clear perivascular mitochondrial clustering together with the marked reduction in mitochondria-ER contact sites in reactive Mfn2^cKO^ astrocytes provides an opportunity for investigating potential functional consequences confined to this cellular compartment. Interestingly, protein expression levels belonging to a Calcium Transport pathway in our IPA analysis were selectively upregulated in Mfn2^cKO^ astrocytes (Figure S3F). In particular, analysis of proteins known to regulate Ca^2+^ influx/efflux through the mitochondrial, ER and plasma membranes revealed differential yet pronounced changes, with a clear trend towards elevated expression of Ca^2+^ channels and transporters in mitochondrial as well as plasma membranes specifically in Mfn2^cKO^ astrocytes (Figure 4K). We thus focused our analysis on local astrocytic Ca^2+^ dynamics (Volterra et al., 2014), as mitochondria-ER tethering domains are known to play a major role in mediating mitochondrial Ca^2+^ uptake and by consequence in regulating cytosolic Ca^2+^ handling mechanisms (Csordas et al., 2018; Rizzuto et al., 2012).

We first evaluated the extent of mitochondrial Ca^2+^ uptake by stereotactically delivering an astrocyte-specific AAV expressing the calcium indicator GCaMP6f targeted to the mitochondrial matrix (mitoGCaMP6) into the cerebral cortex of Mfn2^cKO^ or control littermates, and concurrently inflicted a unilateral SW lesion in the injected area (Figure 5A). We then conducted 2-photon laser scanning microscopy (2PLSM) at 7 or 28 days after SW in freshly prepared brain slices. Imaging was carried out in sessions of 3 minutes each, which corresponded to a time window during which mitochondrial movement or fusion-fission dynamics - as examined via photoactivatable mito-GFP experiments in comparable settings - were negligible (Figure S5A-C), thus allowing a reliable quantification of local relative changes in mitoGCaMP6 signal. We also developed a dedicated algorithm (which we termed AstroSparks, see methods) permitting a semi-automated identification and quantification of spontaneous mitochondrial Ca^2+^ transients, including their activity, frequency, amplitude and duration (Figure 5B). This allowed us to reveal that, in resting astrocytes, perivascular mitochondria are intrinsically more active but display a lower amplitude in their Ca^2+^ transients than mitochondria localized in branches and branchlets (Figure 5C-D). Analysis of Mfn2^cKO^ astrocytes under uninjured conditions (Figure 5E) disclosed an intrinsically lower mitochondrial Ca^2+^ activity within their end-feet (46.6% active ROIs of all ROIs per cell) as compared to Mfn2^WT^ astrocytes (68.2 % active ROIs of all ROIs per cell) (Figure 5F). Interestingly, following SW-injury Mfn2^WT^ astrocytes displayed a peculiar pattern in their mitochondrial Ca^2+^ uptake dynamics that mirrored the morphological changes in mitochondrial network architecture described in Figure 2E: by 7 days (i.e. the peak of mitochondrial fragmentation) the extent of active mitochondria was visibly reduced (55.3 % of all ROIs per cell), whereas by 28 days (the time when mitochondrial tubular morphology had been re-established) this percentage had reverted to levels comparable to uninjured conditions (62.8 %) (Figure 5F). Likewise, most of the other parameters pertaining to Ca^2+^ uptake dynamics, particularly the frequency and duration of Ca^2+^ events per mitochondrion, also followed a reversible pattern over time in Mfn2^WT^ astrocytes (Figure 5G). In contrast, analysis of Mfn2^cKO^ astrocytes revealed that mitochondria in these cells are virtually unresponsive to injury-induced changes of mitochondrial Ca^2+^ uptake all through the analysed times (Figure 5F-G). In particular, the values of frequency, amplitude and duration of Ca^2+^ transients were not only already affected in absence of any SW-injury, but also compared rather well with the 7-day time-point of the Mfn2^WT^ group (Figure 5G), suggesting that primary alterations in mitochondrial network morphology (i.e., towards fragmentation) and mitochondria-ER tethering *per se* are, at least in part, responsible for the changes in mitochondrial Ca^2+^ uptake observed here.

**Figure 5.**
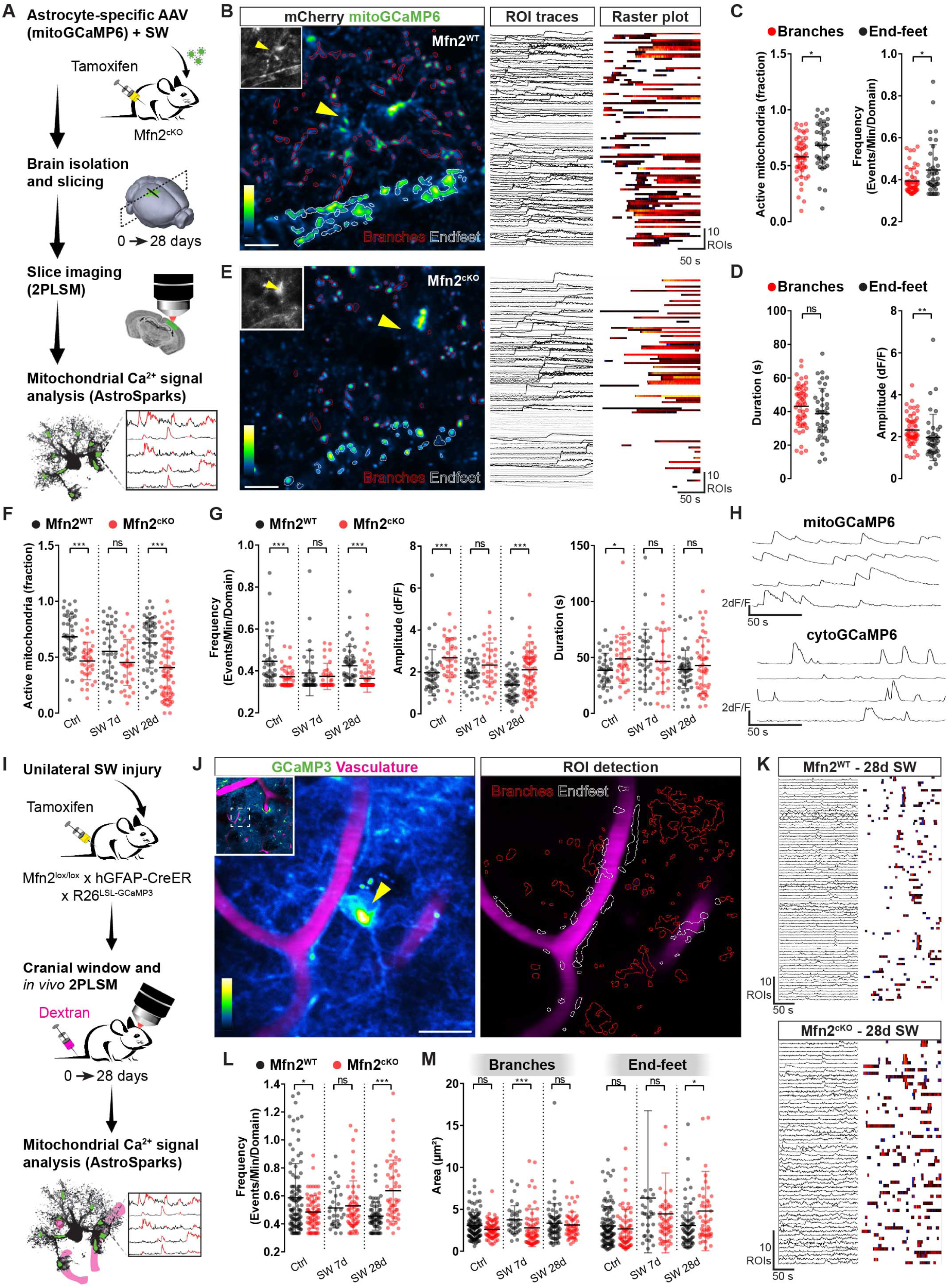
Compromised mitochondrial Ca^2+^ uptake dynamics and abnormal cytosolic Ca^2+^ activity in Mfn2^cKO^ astrocytic end-feet. **(A)** Schematic showing mitoGCaMP6f expression in astrocytes followed by 2PLSM and subsequent AstroSparks analysis. **(B)** Example of a mitoGCaMP6-expressing Mfn2^WT^ astrocyte in brain slice following AstroSparks processing and ROI detection (ROIs in end-feet are depicted in white, branches are in red; soma was excluded). Inset displays cytosolic mCherry (co-expressed with mitoGCaMP6), utilized to identify the end-feet. Bar, 10 µm. Right panels depicts individual ROI traces and the corresponding raster plot. **(C-D)** Quantification of mitochondrial Ca^2+^ transients in branches and end-feet of uninjured Mfn2^WT^ astrocytes (n= 41-53 cells collected from 3 mice). **(E)** Example of a mitoGCaMP6-expressing Mfn2^cKO^ astrocyte with corresponding ROI traces and raster plot. Bar, 10 µm. **(F)** Quantification of active mitochondria in Mfn2^WT^ (n= 40-56 cells, 3 mice/condition) and Mfn2^cKO^ (n= 36-73 cells, 2-3 mice/condition) astrocytic end-feet. **(G)** Quantification of frequency, amplitude and duration of mitochondrial Ca^2+^ transients of the astrocytes shown in **F**. **(H)** Example of mitochondrial and cytosolic Ca^2+^ traces. **(I)** Experimental setting utilized for Ca^2+^ imaging of Mfn2^cKO^ astrocytes *in vivo*. **(J)** Example of a GCaMP3-expressing astrocyte imaged *in vivo* following ROIs detection (excluding the soma). Bar, 20 µm. **(K)** Examples of ROI traces and corresponding raster plots in Mfn2^WT^ and Mfn2^cKO^ astrocytes at 28 days post-SW. **(L)** Average frequency (end-feet) and **(M)** area of Ca^2+^ transients quantified in Mfn2^WT^ (n= 35-111 cells, 2-3 mice/condition) and Mfn2^cKO^ astrocytes (n= 51-73 cells, 2-3 mice/condition). *, p < 0.05, **, p < 0.01, ***, p < 0.001 (nonparametric Mann-Whitney t-test). See also Figure S5.

Analysis of slices containing cytoGCaMP6-expressing astrocytes revealed plain differences with regard to Ca^2+^ transients taking place in the cytosol as compared to mitochondria (Figure 5H and S5D-E). In particular, cytosolic transients in uninjured Mfn2^WT^ astrocytes were markedly shorter in duration and, on average, higher in frequency than mitochondrial ones (Figure 5H and S5E-F), consistent with a role played by mitochondria in rapidly buffering Ca^2+^ ions following cytosolic influx (Rizzuto et al., 2012). SW-injury in cytoGCaMP6-expressing Mfn2^WT^ astrocytes significantly modified perivascular cytosolic transients at 7 days (Figure S5F), yet these changes were not fully reversed by 28 days post-SW, suggesting the emergence of long-lasting alterations in the expression of membrane Ca^2+^ transporters and/or handling mechanisms that may persist up to 1 month after injury. Notably, the frequency of cytosolic transients was significantly altered in resting astrocytes upon conditional deletion of *Mfn2*, but not *Mfn1* (Figure S5E-F), and culminated in an exaggerated Ca^2+^ activity (i.e., frequency and amplitude of events) by 28 days post-SW (Figure S5F), thus validating our Mfn2^cKO^ proteomic dataset (Figure 4C). Interestingly, similar changes in Ca^2+^ activity were also observed in astrocyte branches (Figure S5G), suggesting that lack of MFN2 affected mitochondrial and cytosolic Ca^2+^ frequency dynamics to an overall comparable extent in all astrocytic territories.

While slice imaging allowed us to identify the overall changes in astrocytic mitochondrial and cytosolic Ca^2+^ activity following SW-injury, it precluded the possibility to examine in detail the regionalized Ca^2+^ dynamics within an intact neurovascular unit. To circumvent this caveat, we performed 2PLSM of Mfn2^cKO^ astrocytes in anesthetized animals *in vivo* following cranial window implantation and concurrent vasculature labelling with dextran-red (Figure 5I). For these experiments we introduced the inducible reporter line GCaMP3 floxed-stop (Zariwala et al., 2012) in our Mfn2^cKO^ mice, thus allowing for a systematic analysis of subcellular changes in cytosolic Ca^2+^ activity without the need to inject any AAV. In particular, dextran labelling allowed us to unambiguously identify perivascular end-feet in GCaMP3-expressing astrocytes *in vivo*, and by exclusion the main branches and branchlets (Figure 5J). Analysis of Ca^2+^ frequency in this setting confirmed that Mfn2^WT^ astrocytes undergo substantial alterations in response to SW-injury peaking at 7 days and persisting up to 28 days (Figure 5L and S5G). Importantly, by this time *Mfn2* deletion led to an abnormal frequency of Ca^2+^ events which resulted in significantly higher rates of perivascular transients (0.64±0.03 events/min/domain in Mfn2^cKO^ astrocytes vs 0.46±0.01 events/min/domain in controls) (Figure 5K-L). While this phenotype was present both in end-feet and branches (Figure S5G), analysis of the spatial spreading of Ca^2+^ transients within astrocytic territories revealed that prominent and enduring changes (i.e. broader transients) up to 28 days post-SW were a unique feature of perivascular compartments in astrocytes lacking MFN2 (average transient size of 4.98±0.67 μm^2^ in Mfn2^cKO^ vs 3.25±0.33 μm^2^ in Mfn2^WT^) (Figure 5M). This hallmark was masked at 7 days post-SW, when control astrocytes also showed broader transients presumably due to their conspicuous mitochondrial fragmentation and reduced mitochondrial Ca^2+^ uptake (Figure 5G), yet this specificity for the end-feet indicates that injury-induced accumulation of mitochondria-ER contact sites at this location helps to demarcate a region of distinctive Ca^2+^ signalling and, supposedly, metabolic supply which may potentially contribute to vascular remodelling following injury.

### Astrocyte mitochondrial fusion dynamics are required for vascular remodeling following injury

In order to understand if the observed structural and functional changes in perivascular mitochondrial-ER contact sites exhibited by Mfn2^cKO^ astrocytes may have any direct consequence for vascular remodelling, we performed a systematic analysis of the vascular plexus following cortical injury. SW-injured mice were intravenously infused with dextran-red shortly before sacrifice and their cortices processed for clearing and 2PLSM to obtain a complete overview of the vascular network architecture (Figure 6A). Top views of the first 600μm deep into the cortex revealed that uninjured hemispheres were virtually undistinguishable among Mfn2^cKO^ and Mfn2^WT^ mice, showing comparable density and integrity of the labelled vasculature (Figure 6B). By 7 days post-SW, however, a clear lesion track and a reduction in vascular density became visible in Mfn2^WT^ mice, yet Mfn2^cKO^ cortices showed a much prominent rarefication of the vasculature within the lesion core (Figure 6B). By 28 days, the observed rarefication in control mice appeared virtually resolved at the location where the previous injury had been inflicted. In contrast, the lesion core in Mfn2^cKO^ mice retained much of the alterations identified at 7 days, suggesting an impairment in vascular remodelling following injury (Figure 6B). To quantify the extent of vascularization in the injured area, we optimized a filament tracing analysis utilizing dextran labelling as a mask signal for our volumetric reconstructions (Figure 6C and S6A; see methods) and performed a post-SW time-course analysis in Mfn2^WT^ and Mfn2^cKO^ mice. At the earliest analysed time (3 days), we identified a similar reduction in the density of branch points as well as total length and fractional vascular volume of the network immediately surrounding the injury track (i.e., within a fixed total volume of about 0.2 mm^3^) as compared to uninjured conditions (Figure 6D). Yet, while by 7 days the Mfn2^WT^ group started showing a progressive recovery of these parameters (in particular branch points) which became conspicuous by 28 days, Mfn2^cKO^ injured cortices failed in undergoing significant improvements (Figure 6D). Interestingly, analysis of injured Mfn1^cKO^ mice did not reveal striking dissimilarities compared to wild-type injured mice for any of the examined parameters (Figure S6B-C), in line with the fact that disruption of MFN1 expression alone did not prevent perivascular clustering of mitochondria (Figure 4K-L).

**Figure 6.**
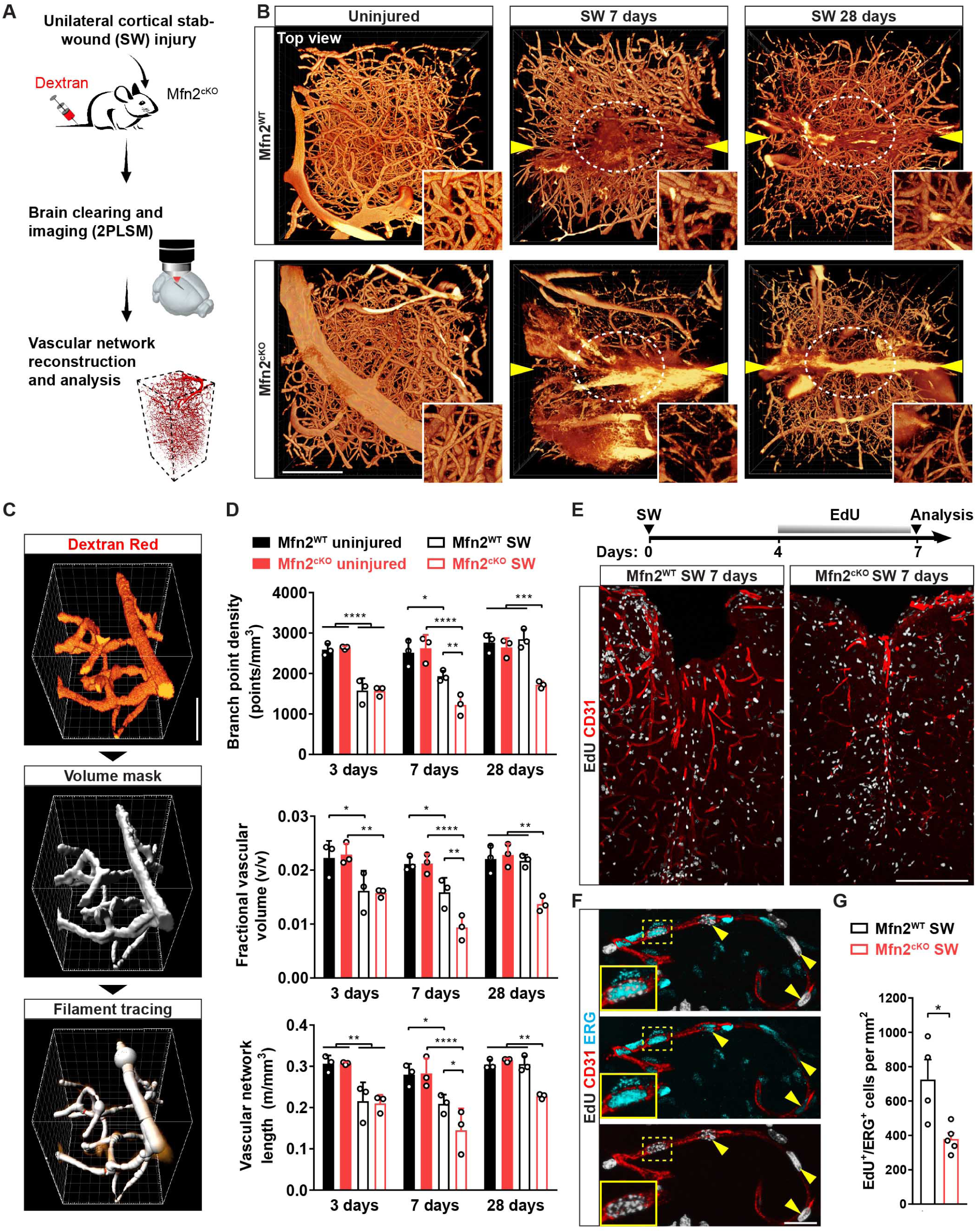
Disruption of astrocyte mitochondrial fusion dynamics impairs injury-induced angiogenesis and vascular remodelling. **(A)** Experimental protocol used for examining the vascular network following injury. **(B)** Top views of the vascular network in reconstructed portions of Mfn2^WT^ and Mfn2^cKO^ cortices. Arrowheads point to the lesion tracks (penetrating lesion perpendicular to the field of view). Insets depict zooms of the lesioned core region (circled in white). Bar, 200 µm. **(C)** Pipeline used for vasculature quantification via filament tracing of the dextran signal. Bar, 30 µm. **(D)** Quantification of branch points, fractional volume and total length of the vascular network in Mfn2^WT^ and Mfn2^cKO^ cortices (n= 3 mice/condition; two-way ANOVA followed by Tukey’s post-hoc test). **(E)** Scheme showing the experimental timeline used for EdU labeling of proliferating cells in SW-injured animals. Lower pictures depict large views of the injured cortex in Mfn2^WT^ and Mfn2^cKO^ mice at 7 days following immunostaining for CD31 as well as EdU. Bar, 200 µm. **(F)** Magnification of a microvessel proximal to the lesion track showing the presence of endothelial cells (CD31+/ERG+) that have incorporated EdU during the previous 3 days. **(G)** Quantification of proliferating ERG+ cells within the area surrounding the lesion track in Mfn2^WT^ and Mfn2^cKO^ mice at 7 days post-SW (n= 4-5 mice/condition; nonparametric Mann-Whitney t-test). *, p < 0.05, **, p < 0.01, ***, p < 0.001. See also Figure S6 and S7.

To gain insights into the mechanisms underlying the impaired vascular remodelling of Mfn2^cKO^ mice, we examined their angiogenic response to injury by 7 days post-SW. Proliferating cells were quantified by supplying EdU to mice during the last 3 days before sacrifice (Figure 6E), and labelled cells examined for their positivity to the ETS-transcription factor ERG, an endothelial marker known to promote angiogenesis (Birdsey et al., 2008). Inspection of brain sections in proximity to the lesion track in Mfn2^WT^ mice revealed a number of EdU+/ERG+ cells along CD31+ vessels, indicative of neoformed vessels containing endothelial cells that did proliferate during this time (Figure 6F). Importantly, in Mfn2^cKO^ mice the overall extent of CD31+ vessels alongside the density of EdU+/ERG+ cells appeared markedly reduced (Figure 6E and G), despite comparable numbers of total proliferating EdU+ cells as well as SOX2+ astrocytes to Mfn2^WT^ mice (Figure S6D-F). These data were corroborated by examining the total density of astrocytes in brain sections derived from injured Mfn2^cKO^ mice and control littermates expressing a Cre-dependent cytosolic tdTomato reporter (Figure S7A). Importantly, quantification of astrocyte numbers and inspection of perivascular end-feet in these tdTomato-expressing Mfn2^cKO^ mice revealed no overt changes as compared to Mfn2^WT^ mice (Figure S7A-C), ruling out possible effects due to astrocyte degeneration or prominent changes in their proliferative capacity.

Together, these results indicate that lack of MFN2 in reactive astrocytes compromises vascular remodelling after injury by limiting angiogenesis, while astrocyte proliferation as well as general morphology appear preserved.

### Forced enrichment of mitochondria-ER contact sites in perivascular end-feet rescues vasculature remodeling in absence of mitochondrial fusion

While lack of MFN2 in reactive astrocytes is sufficient to impair the formation of new vessels after SW-injury, it still remains unclear whether this effect is mediated by defective mitochondrial fusion *per se* rather than by disrupted mitochondria-ER tethering. We thus asked whether perivascular enrichment of mitochondria-ER contact sites may be sufficient to re-establish vascular remodeling after injury. To address this question, we took advantage of a previously validated strategy to forcefully anchor mitochondria to ER membranes using a genetically-encoded synthetic linker (Csordas et al., 2006) that we expressed in Mfn2^cKO^ mice via an astrocyte-specific AAV (pAAV-hGfaABC_1_D-OMM-mRFP-ER) (Figure 7A). We reasoned that, since the overall extent of perivascular ER tubules appeared to large degree conserved in absence of MFN2 prior and after injury (Figure 3C-D and S4D-E), anchoring mitochondria to ER tubules before tamoxifen treatment (and *Mfn2* deletion) by means of this irreversible linker may enhance the extent of contact sites irrespective of subsequent changes in morphology and fission-fusion dynamics. The construct encoding for this linker contains a monomeric RFP fused on one side to the outer mitochondrial membrane (OMM)-targeting sequence of mAKAP1 and on the other to the ER membrane-targeting sequence of yUBC6 (Figure 7A) (Csordas et al., 2006). Its expression in different cellular systems markedly expands the interface area between mitochondria and ER, resulting in mRFP labelling of the OMM (Arruda et al., 2014; Csordas et al., 2006; Csordas et al., 2010). Few weeks after intracortical delivery of this AAV-linker (or its AAV control lacking the ER targeting sequence), mice were treated with tamoxifen to induce *Mfn2* deletion (Figure 7A) followed by SW-injury and mitochondrial network analysis. At the single-astrocyte level, the overall morphology of the mitochondrial network was not significantly rescued, with most mitochondria still appearing visibly fragmented even in absence of injury (Figure 7B), as expected in astrocytes lacking mitochondrial fusion. However, we noticed that the amount of mRFP+ mitochondria decorating vessel cross-sections was visibly increased in astrocytes transduced with the AAV-linker as compared to controls in both resting and injured conditions (Figure 7B-D). Importantly, this effect was independent of mitochondrial morphological changes within the end-feet, as AAV-linker expression was not able to restore tubular mitochondria (Figure S7D). To understand if this manipulation also functionally modified the microenvironment of perivascular end-feet, we introduced a cassette encoding for mitoGCamp6f in the OMM-mRFP-ER construct and performed Ca^2+^ imaging in brain slices following AAV cortical delivery *in vivo* (Figure S7E-F). Analysis of mitoGCamp6f in resting Mfn2^cKO^ astrocytes revealed that AAV-linker transduction modified the extent of mitochondrial Ca^2+^ uptake by increasing both the percentage of active mitochondria and their frequency dynamics (Figure 7E and Figure S7G-H) to levels almost comparable to Mfn2^WT^ astrocytes (Figure 5F-G), indicating that this forced tethering was sufficient to enhance mitochondrial Ca^2+^ uptake in absence of MFN2.

**Figure 7.**
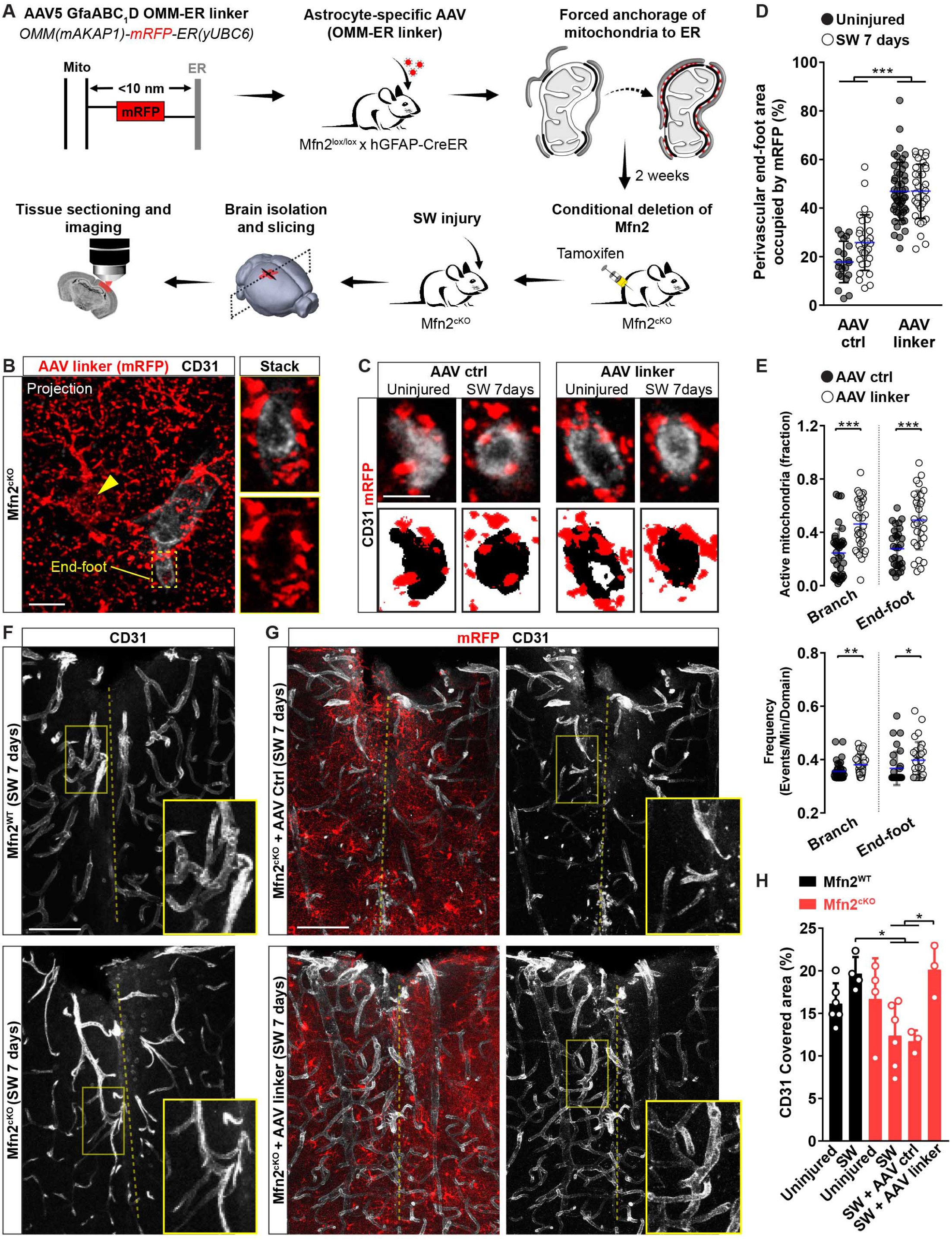
Forced enrichment of mitochondria-ER tethers in perivascular end-feet rescues vascular remodeling in injured Mfn2^cKO^ mice. **(A)** Experimental plan for expressing the artificial mitochondria-ER linker (OMM-mRFP-ER) in Mfn2^cKO^ astrocyte *in vivo*. **(B)** Example of an AAV-linker-expressing Mfn2^cKO^ astrocyte (arrowhead points to the soma) showing mRFP-labelled mitochondria and a nearby vessel. Zooms depict the vessel cross-section. Bar, 10 µm. **(C)** Examples of vessel cross-sections following expression of the AAV-ctrl or AAV-linker in Mfn2^cKO^ astrocytes. Bar, 5 µm. **(D)** Quantification of perivascular mRFP+ mitochondrial density displayed as area fraction (n≥ 21 vessel sections; one-way ANOVA followed by Tukey’s post-hoc test). **(E)** Quantification of mitochondrial Ca^2+^ uptake in Mfn2^cKO^ astrocytes following expression of mitoGCaMP6f in the AAV-ctrl and AAV-linker (n≥ 30 cells; nonparametric Mann-Whitney t-test). **(F)** Examples of vasculature density (CD31+) in Mfn2^WT^ and Mfn2^cKO^ sections at 7 days post-SW (dashed line points to the lesion track). Bar, 80 µm. **(G)** Examples of vasculature density in injured Mfn2^cKO^ cortices transduced with the AAV-Ctrl or AAV-linker. Bar, 80 µm. **(H)** CD31 area fraction in Mfn2^WT^ and Mfn2^cKO^ cortical sections under the indicated conditions (n≥ 3 mice/condition; one-way ANOVA followed by Tukey’s post-hoc test). *, p < 0.05, **, p < 0.01, ***, p < 0.001. See also Figure S7.

We next analysed the extent of vasculature remodelling induced by SW-injury in the area subjected to AAV transduction. Visual inspection of CD31 immunoreactivity confirmed that non-transduced Mfn2^cKO^ cortices were characterized by a less elaborated vascular network in the injured area as compared to Mfn2^WT^ cortices (Figure 7F and H). Importantly, while injection of the AAV-ctrl did not overtly change the extent of CD31+ vessels by 7 days post-SW in Mfn2^cKO^ mice, AAV-linker expression significantly enhanced vascular complexity to levels almost indistinguishable from those of Mfn2^WT^ mice (Figure 7G and H). Accordingly, AAV-linker expression increased the number of branch points and total vascular length in Mfn2^cKO^ cortices as compared to AAV-ctrl expression (Figure S7I). Together, these results indicate that, within the examined time frames, forced enrichment of mitochondrial-ER tethering in Mfn2^cKO^ astrocytic perivascular end-feet is sufficient to restore vascular remodelling following injury.

## Discussion

We have shown that a profound reorganization of the mitochondrial network in astrocytes responding to acute injury underlies their ability to create a spatially defined mitochondrial-enriched domain in perivascular end-feet. Astroglial end-feet appear to be naturally enriched in elaborated mitochondrial morphologies and bundles of ER tubules, which is in line with recent observations (Mathiisen et al., 2010; Moss et al., 2016), yet during the first week that follows injury these cellular sites experience a further accumulation of mitochondria as a result of coordinated fusion-fission dynamics. While mitochondrial biogenesis or trafficking are also likely to contribute in this process, mitochondrial fusion in particular was required to promote the formation of this localized clustering as deletion of *Mfn2* not only prevented this response, but also significantly altered the extent of contact sites with the ER, thus affecting local Ca^2+^ dynamics. Importantly, the extent of mitochondrial accumulation and ER tethering in astrocytic end-feet had direct consequences for microvasculature remodeling: while depletion of mitochondrial-ER contact sites impaired angiogenesis and vascular complexity in lesioned cortices, mitochondrial and ER-tether enrichment had opposite results and rescued vascular density even in absence of mitochondrial fusion. This finding is reminiscent of an equally enhanced accumulation of mitochondria to new axonal sprouts following axotomy experiments, a process which has implications for axon regeneration (Han et al., 2016; Mar et al., 2014; Misgeld et al., 2007). Along this line, our data support the notion that enrichment of mitochondria and mitochondria-ER contact sites in astrocytic end-feet does not simply identify a general trait of cellular reactivity but rather a mechanism that is triggered to ensure the formation of an active metabolic compartment with direct implications for vascular remodelling.

Our experiments performed on astrocyte-specific Mfn2^cKO^ mice were specifically designed to manipulate the mitochondrial network shortly (∼2 weeks) before inflicting the SW-injury, thereby allowing mice to develop and reach adulthood with normal MFN2 expression until the first day of tamoxifen treatment. This time was sufficient to elicit a significant drop in MFN2 protein expression *in vivo*, which was mirrored by evident changes in mitochondrial morphology and ultrastructure. This indicates that mitofusins have a relatively rapid turnover in astrocytes and allowed us to focus on the acute effects resulting from lack of mitochondrial fusion. While this may explain the seemingly intact cristae morphology observed in Mfn2^cKO^ astrocytes, in contrast to developmental knockout studies (Chen et al., 2007; Lee et al., 2012), it is of particular interest the fact that morphological changes towards circular and fragmented mitochondria were accompanied by a clear reduction in the extent of MAM domains with the ER in astrocytic end-feet. In cell lines, MFN2 has been repeatedly reported to regulate the extent of tethering between these two organelles, with a pro-tethering (de Brito and Scorrano, 2008; Naon et al., 2016) rather than an anti-tethering activity (Filadi et al., 2015) being validated also in other *in vivo* studies (Luchsinger et al., 2016; Schneeberger et al., 2013). Here, a reduction in MAMs and an increased mitochondria-ER distance in astrocytes supports a similar pro-tethering role of MFN2. However, we cannot entirely exclude that these may partly develop as secondary effects due to morphological changes of the mitochondrial network in perivascular end-feet.

One of the key findings of our study is the observation that the enrichment of mitochondria-ER tethering within end-feet contributes to regulate the local environment surrounding microvessels *in vivo* and *ex vivo*, as revealed by Ca^2+^-imaging experiments performed with the GCamp6f sensor. Interestingly, our analysis revealed that the end-foot is characterized by distinctive mitochondrial Ca^2+^ uptake dynamics when compared to peripheral branches of the same cell, which may be justified by the enrichment in mitochondria-ER contact sites precisely in perivascular processes. Astrocytes possess a remarkably complex Ca^2+^ activity on account of their highly ramified morphology (Bindocci et al., 2017; Shigetomi et al., 2016) and changes in the pattern of spontaneous and stimulus-induced Ca^2+^ transients have been shown to associate with synaptic transmission and vascular tone (Bindocci et al., 2017; Tran et al., 2018; Wang et al., 2006). Besides the ER, mitochondria are also well known for being integral components of Ca^2+^ signalling in cells given their significant Ca^2+^ buffering capacity which is primarily regulated by the mitochondrial calcium uniporter (MCU) complex (Baughman et al., 2011; De Stefani et al., 2011). Calcium uptake can potentially modify mitochondrial bioenergetics (Giorgi et al., 2018), but also the magnitude and spread of cytosolic Ca^2+^ transients and thus have important effects on key signalling events in cells, including astrocytes *in vitro* (Jackson and Robinson, 2015; Li et al., 2014; O’Donnell et al., 2016; Parnis et al., 2013; Parpura et al., 2011; Reyes and Parpura, 2008; Stephen et al., 2015) and *in vivo* (Agarwal et al., 2017). As MCU exhibits low Ca^2+^affinity, mitochondrial Ca^2+^ influx predominantly occurs at sites of elevated Ca^2+^ concentrations, i.e. mitochondria-plasma membrane and mitochondria-ER tethering domains (Hayashi et al., 2009; Rizzuto et al., 2012). Intriguingly, manipulation of MFN2 expression levels has been shown to alter mitochondrial Ca^2+^ buffering capacity in cells as a consequence of its regulatory role on mitochondria-ER tethering domains (de Brito and Scorrano, 2008; Filadi et al., 2015; Luchsinger et al., 2016; Naon et al., 2016). Consistent with these earlier reports, here we find that a marked dampening of mitochondrial Ca^2+^ activity in Mfn2^cKO^ astrocytes *in situ* renders these cells virtually insensitive to the changes in Ca^2+^ dynamics induced by injury, while forced expression of a mitochondria-ER synthetic linker alone is sufficient to restore Ca^2+^ uptake even in absence of mitochondrial fusion, thus indicating that tethering domains play a major role in astrocyte Ca^2+^ handling mechanisms. Importantly, while this impaired mitochondrial Ca^2+^ uptake is likely to impact local perivascular bioenergetics in Mfn2^cKO^ astrocytes, it certainly translates into long-term alterations in cytosolic Ca^2+^ activity which, at the level of the end-feet, manifest as Ca^2+^ transients wider and more frequent than those observed in control end-feet. It is thus tempting to speculate that these abnormal Ca^2+^ transients may affect astrocytic perivascular function, including secretion of vasoactive or pro-angiogenic molecules, yet the exact consequences of this altered Ca^2+^ activity for vascular remodelling remains to be clarified. In future studies, it will thus be interesting to assess if mitochondrial Ca^2+^ uptake blockade, for instance via astrocyte-specific genetic manipulation of MCU, may also affect vascular remodelling in injury settings.

Unexpectedly, we found that abrogation of astrocyte MFN2 and the ensuing disruption of mitochondria-ER contact sites was sufficient to impair angiogenesis and vascular remodeling after injury. While we propose this effect to be primarily mediated by a faulty metabolic domain at the gliovascular interface, at this stage we can only argue what the exact signalling might be that facilitates a vascular response in physiological conditions. Interestingly, MFN2-mediated signalling has been implicated in regulating cell proliferation cell-autonomously in vascular smooth muscle cells (Chen et al., 2004), however here we did not find overt changes in astrocyte proliferation or survival in our system. Also, the fact that astrocytes can sustain a glycolytic metabolism for extended periods of time (Supplie et al., 2017) argues against a primary role of OXPHOS in this regard. In line with this notion, and despite a partial alteration on OXPHOS components identified in our proteomics of Mfn2^cKO^ astrocytes, we were unable to reveal major changes in TCA cycle metabolites or amino acid biosynthesis following ^13^C_6_-Glucose administration *in vivo*. While these data do not completely rule out potential local changes in energy metabolism restricted to the perivascular end-feet, mitochondrial cristae ultrastructure appeared intact in Mfn2^cKO^ reactive astrocytes even at late time points after injury, thus indicating that mitochondrial metabolism *per se* may not be strongly affected in our model. Moreover, forced induction of mitochondria-ER tethering domains via AAV-linker expression alone in Mfn2^cKO^ mice was sufficient to restore vascular remodelling, providing additional evidence for the presence of still functional mitochondria. Thus, one intriguing possibility is that this close apposition of a dense supply of mitochondria-ER contact sites at the vascular interface may favour either the local accumulation of specific signalling molecules or contribute to generate locally a chronic metabolic environment (Al-Mehdi et al., 2012; Booth et al., 2016; Lopez-Fabuel et al., 2016), which may act non cell-autonomously in assisting the angiogenic response during the days that follow the initial insult (Wong et al., 2017). Alternatively, a steady and local supply of key astrocytic biosynthetic intermediates, as those generated by the TCA cycle (Lovatt et al., 2007), or ATP itself may contribute to keep fuelling the remodelling of the gliovascular interface (Boulay et al., 2017; Rangaraju et al., 2019) as well as restore perivascular barrier (Voskuhl et al., 2009) or clearance functions, in particular of toxic metabolic by-products (Iliff et al., 2012). Ultimately, a combination of multiple factors, possibly converging onto the localized release of pro-angiogenic signalling molecules (Sweeney et al., 2016), are likely to participate in regulating astrocyte-mediated vascular remodelling following injury.

In conclusion, our study provides insights into the changes in mitochondrial structure and function experienced by astrocytes during their response to cerebrovascular damage, but also it identifies an important mechanism through which these cells directly contribute to vascular remodelling in the injured brain. Successful molecular dissection of the precise metabolic pathways playing a role in this process may therefore hold promise for therapeutic interventions to ameliorate tissue repair.

## Author Contributions

J.G. performed and analysed most of the experiments. E.E. contributed to MACS and FACS experiments. P.P. developed the custom algorithm for Ca^2+^ analysis. V.S. and M.J. generated and validated AAVs. H.M.J. and A.S. contributed to experiments. K.F.D. and C.K. supported with cell sorting. A.G. and K.K.C. provided reagents. C.F. performed mass-spec and initial analysis. P.G. performed metabolomics and initial analysis. E.M. contributed to proteomic and metabolomics analysis. J.G., E.M. and M.B. prepared figures. E.M. and M.B. developed the concept, designed experiments, analysed data and wrote the paper. All authors revised the manuscript.

## Acknowledgements

We thank N.G. Larsson for providing mitoYFP floxed-stop, Mfn1 floxed and Mfn2 floxed mice. M. Götz and F. Kirchhoff for Glast::CreERT2 mice. L. Uhrbom and E. Holland for hGFAP-TVA mice. T. Langer for insightful comments. E.L. Snapp for ER-targeted probes. G. Hajnoczky for synthetic linkers. N. Toni for advices on EM. I. Atanassov for support with IPA analysis. B. Fernando, T. Öztürk and S. Perin for excellent technical assistance. J. Matutat, G. Piper, D. Schneider and the other members of the CECAD in vivo facility for excellent assistance. G. Wani, S. Wendler, K. Ndoci and T. Eriksson for general lab support and discussions. All members of the CECAD imaging facility for assistance with microscopes and software. S. Müller and the team of the CECAD proteomics core facility for technical assistance. This work was supported by the Deutsche Forschungsgemeinschaft (SFB1218 A07), European Research Council (ERC-StG-2015, grant number 67844), Köln Fortune and UoC Advanced Postdoc Grant to M.B; the Deutsche Forschungsgemeinschaft (SFB1218 Z03) to A.S.; and the Deutsche Forschungsgemeinschaft (SFB870) to K.K.C. E.M. is recipient of an Advanced Postdoc Grant (Deutsche Forschungsgemeinschaft, SFB1218). C.K. acknowledges the ISAC SRL Emerging Leaders Program.

## Author Information

The authors declare no competing financial interests.

## Legends to Supplemental Figures

**Figure S1. Related to Figure 1. Characterization of organelle distribution across astrocytic territories *in vivo*. (A)** Left panel: example (confocal z-projection) of a cortical astrocyte transduced with an astrocyte-specific virus encoding for mitoRFP. The location of one astrocytic end-foot and soma (S) are depicted. Right panel: rendered image of mitoRFP (following deconvolution) of a single stack and the corresponding immunoreactivity for CD31 of nearby microvessels. Low panels shows zooms of each respective yellow boxed region. Bars, 15 µm. **(B)** Left panel: example (confocal z-projection) of a cortical astrocyte transduced with an astrocyte-specific virus encoding for ER-GFP. The location of astrocytic end-feet and soma (S) are depicted. Right panel: rendered image of ER-GFP (following deconvolution) of a single stack and the corresponding immunoreactivity for CD31 of nearby microvessels. Low panels shows zooms of each respective yellow boxed region. Bars, 15 µm. **(C)** Example of a cortical astrocyte transduced with an astrocyte-specific AAV encoding for a lysosomal marker (Emerald-Lamp1) and cytosolic mCherry. The location of astrocytic territories including end-foot, branches/branchlets and soma (S) is depicted. Immunostaining for CD31 shows the presence of nearby microvessels. The right panel shows the Emerald-Lamp1 channel reporting on the distribution of lysosomes. Bar, 15 µm. **(D)** Zooms (surface rendered) of the boxed areas shown in **C**. Bar, 5 µm. **(E)** Example of a cortical astrocyte transduced with an astrocyte-specific AAV encoding for a peroxisomal marker (mCherry-Perox) and cytosolic BFP. The location of astrocytic territories including end-foot, branches/branchlets and soma (S) is depicted. Immunostaining for CD31 shows the presence of nearby microvessels. The right panel shows the mCherry-Perox channel reporting on the distribution of peroxisomes. Bar, 15 µm. **(F)** Zooms (surface rendered) of the boxed areas shown in **E**. Bar, 5 µm. **(G)** Example of a portion of cortex in a brain section from tamoxifen-induced Glast::CreER^T2^ x R26^LSL-tdTomato^ mice immunostained for the endothelial marker CD31, showing the extent of perivascular end-feet wrapping around the vasculature. Bar, 50 µm. **(H)** EM picture of a similar specimen as in **G** following immuno-gold processing against RFP. A superimposed red shadow identifies the location of the perivascular end-foot enriched in gold particles. The zoom on the right illustrates the localization of gold particles within the end-foot surrounding the basal lamina.

**Figure S2. Related to Figure 2. Remodeling of astrocyte mitochondrial and ER networks following SW-injury. (A)** Example of brain section from tamoxifen-induced hGFAP::CreER^TM^ x R26^LSL-mitoYFP^ mice immunostained for the astrocytic marker S100β. Left inset: zoom of a single S100β+/mitoYFP+ astrocyte. Right inset: quantification of recombination efficiency in the cortex. Bars, 20 µm. **(B)** Examples of mitoYFP+ astrocytes in control (uninjured) and injured conditions (SW 7 days, astrocyte proximal to the lesion track) showing the presence of CD45+ leukocytes (labeled in cyan) after SW injury. Bars, 15 µm. **(C)** Examples of a mitoYFP+ (left) and an ER-GFP+ (right) astrocyte following injury (SW 7 days) showing co-labeling for the endothelial marker CD31. Zooms of the boxed regions depict the end-foot. Bar, 20 µm. **(D)** Quantification of astrocytic mitochondrial mass (total mitoYFP volume per astrocyte) in control (uninjured conditions, time 0) or following stab-wound injury (SW) at 7 and 28 days. Mitochondrial mass was normalized to that of control astrocytes at time 0. N= 22 (time 0), 32 (time 7 days) and 29 (time 28 days) astrocytes obtained from 3 different mice for each time point (one-way ANOVA followed by Kruskal-Wallis test). **(E)** Density of mitoYFP+ signal in peripheral branches of resting or reactive astrocyte at the indicated conditions (n≥ 33 astrocytes obtained from 3 mice/condition) (one-way ANOVA followed by Kruskal-Wallis test). **(F)** Scheme depicting the approach utilized for FACS and proteomic analysis of astrocytes following SW-injury. **(G)** Heat maps of normalized LFQ (label-free quantification) intensities of detected proteins regulating mitochondrial fission and fusion dynamics at the indicated time points after injury and color-coded according to their z-score (n= 6 mice per time point). **(H)** Heat maps of normalized LFQ (label-free quantification) intensities of detected proteins associated to mitochondrial biogenesis (Tfam and Nrf1) as well as with mitochondrial mass (Timm and Tomm proteins) (n= 6 mice per time point). Significant changes (-log_10_ of the *p*-value ≥1.3) are indicated with an asterisk. **(I)** Examples of a reconstructed ER-GFP astrocyte (same as Figure 2J) following volume masking, segmentation of the indicated compartments (end-feet, soma and branches) and subsequent fractionation of the ER-GFP signal in each of these compartments to obtain the signal densities displayed in panel **L**. Bar, 20 µm. **(J)** Quantification of vessel diameter in the same dataset utilized to examine the ER-GFP perivascular g-ratio of Figure 2K. The plot shows data collected during a time course ranging from time 0 (uninjured) to 28 days after injury (n≥ 35 vessels/time-point; nonparametric Kruskal-Wallis test). **(K)** 3D examples of ER-GFP labelled reactive astrocytes at 7 and 28 days post-SW. Volume segmentation into end-feet (according to direct contact with the labelled vasculature), soma and branches is shown in different colors. Lower panels depict the ER-GFP signal density in pseudocolors. A zoom of a perivascular end-foot is shown. Bars, 10 µm. **(L)** Quantification of the fractional ER-GFP signal density across the three indicated astrocytic compartments in uninjured (n= 15 cells, 3 mice) or injured (7 days, n= 13 cells, 3 mice; 28 days, n= 22 cells, 2 mice) astrocytes. **(M)** Quantification of ER-GFP total volume per astrocyte. Right graph: average number of end-feet for the analysed ER-GFP expressing astrocytes as in **L** (one-way ANOVA followed by Dunn’s post-hoc test). *, p < 0.05, **, p < 0.01, ***, p < 0.001.

**Figure S3. Related to Figure 3 and 4. Label-free proteomic analysis of reactive Mfn2^cKO^ and Mfn1^cKO^ astrocytes responding to SW-injury. (A)** Scheme showing the genotyping approach used for validating the conditional knock-out of *Mfn1* and *Mfn2* in cortical astrocytes *in vivo*. **(B)** Genotyping of isolated cortices from tamoxifen-induced Mfn1^cKO^, Mfn2^cKO^ and relative control littermates (Mfn1^WT^ and Mfn2^WT^). The upper gels report on the genotyping protocol to detect wild-type and floxed alleles for each gene, while the lower gels report on the deletion (knock-out) band originating from recombined astrocytes. **(C)** Volcano plot of Mfn2^cKO^ reactive astrocytes (∼3280 detected proteins, ∼2500 quantified) showing their relative expression levels (log_2_ fold change) compared to reactive Mfn2^WT^ (Ctrl) astrocytes obtained from tamoxifen-induced littermates. Proteins with a *p*-value ≤ 0.05 (i.e. ≥ 1.3 on the –log_10_ scale) are considered significant. Proteins annotated in the Mitocarta 2.0 are outlined in red (n= 4 Mfn2^cKO^ mice and 3 Ctrl mice). **(D)** Heat map of normalized LFQ intensities of astrocytic markers of reactivity identified in our proteomics dataset and color-coded according to their z-score. Significant changes (-log_10_ of the *p*-value ≥1.3) are indicated with an asterisk at the beginning of each row. **(E)** Plot showing the increased expression of MFN2 in sorted Mfn1^cKO^ astrocytes at 28 days following injury. The left column reports on the distribution of the whole proteome in Mfn1^cKO^ astrocytes. MFN2 expression is significantly up-regulated under these conditions (**, p value <0.01). **(F)** Ingenuity Pathway Analysis (IPA) of the proteome of Mfn2^cKO^ and Mfn1^cKO^ astrocytes disclosing significantly up-(red) and down-regulated (blue) pathways (bars indicate the -log_10_ of the p-value starting with a minimum cut-off of 1.3). Besides several shared pathways, Mfn2^cKO^-specific up-regulated pathways included Wnt/β-catenin, Insulin Receptor Signaling, Methylmalonyl and 2-oxanobutanoate Degradation and Ca^2+^ Transport. Of the down-regulated pathways, OXPHOS and Regulation of eIF4 and p70S6K Signaling appeared to be specific for Mfn2^cKO^ astrocytes (n= 4 Mfn2^cKO^ mice, 4 Mfn1^cKO^ mice and 3 Ctrl mice). **(G)** Heat maps of normalized LFQ (label-free quantification) intensities of detected OXPHOS complex subunits (complexes I to V) color-coded according to their z-score (n= 4 Mfn2^cKO^ mice, 4 Mfn1^cKO^ mice and 3 Ctrl mice). Significant changes (-log_10_ of the *p*-value ≥1.3) are indicated with an asterisk at the beginning of each row. **(H)** Heat maps of normalized LFQ intensities of detected proteins associated to mitochondrial stress responses color-coded according to their z-score. Significant changes (-log_10_ of the *p*-value ≥1.3) are indicated with an asterisk at the beginning of each row.

**Figure S4. Related to Figure 4. Mitochondrial network changes in astrocytes following deletion of *Mfn2* or *Mfn1*. (A)** Surface rendering examples of mitochondrial morphologies detected in Ctrl, Mfn2^cKO^ and Mfn1^cKO^ resting astrocytes (i.e., uninjured animals). Yellow arrowheads point to the soma. Zooms of the boxed areas depict the predominant network morphology in peripheral processes. Bar, 15 µm. **(B)** Examples of ER-RFP+ resting astrocytes (i.e., in uninjured mice) showing the distribution of the ER in relation to CD31 immunostaining. Yellow arrowheads point to the soma. Zooms of the boxed regions depict the end-feet. Bar, 20 µm. **(C)** Top panels: 3D volume reconstructions showing Ctrl, Mfn2^cKO^ and Mfn1^cKO^ astrocytes (arrowheads point to the soma) surrounding dextran-labelled vessels at 7 days post-SW. Bottom pictures show single-stack views highlighting the extent of perivascular mitochondria for each condition. Bar, 20 µm. **(D)** EM pictures of astrocytic end-feet in Mfn2^WT^ and Mfn2^cKO^ mice at 4 weeks post-SW, showing the extent and morphology of perivascular mitochondria. Images were taken in proximity to the lesion track. Insets depict zooms of mitochondrial cristae. EC: endothelial cell; BL: basal lamina. Bars, 500 nm. **(E)** Details of astrocytic end-feet showing the morphology of ER tubules in Mfn2^WT^ and Mfn2^cKO^ mice. Bar, 500 nm. **(F)** Quantification of the indicated ultrastructural parameters in Mfn2^WT^ (n= 18 vessel cross-sections from 3 mice) and Mfn2^cKO^ perivascular end-feet (n= 24 vessel cross-sections from 3 mice; non-parametric Mann-Whitney t-test). ***, p < 0.001.

**Figure S5. Related to Figure 5. Mitochondrial and cytosolic Ca^2+^ dynamics in Mfn2^cKO^ astrocytes. (A)** Schematic illustrating the experimental protocol used to transduce astrocytes in hGFAP-TVA mice with an EnvA-RABV encoding for the photoactivatable mito-GFP sensor (mito-PA-GFP). Seven days after virus delivery, 2PLSM in fresh brain slices was utilized to assess mitochondrial fusion dynamics in transduced cortical astrocytes. ROI photoactivation in selected astrocytic processes was achieved by laser illumination in the UV range (840 nm at 10% of laser power for 10 seconds), which resulted in bright GFP emission. Time-lapse was performed to follow the fate of the photoactivated mitochondria, which in case of fusion occurring would lead to sudden appearance in the GFP channel of “new” (non-photoactivated) mitochondria, with concomitant dilution of GFP signal intensity in initially photoactivated mitochondria undergoing fusion. **(B)** Example of a mito-PA-GFP-expressing astrocyte surrounded by small and large vessels (indicated by yellow arrows) recognizable by the characteristic mitochondrial outlining into tube-like structures. The laser intensity utilized for GFP detection (920 nm) was slightly increased during preliminary acquisition to identify the morphological appearance of weak PA-GFP-expressing mitochondria along processes and putative end-feet. Boxed areas point to selected ROIs prior photoactivation. Panels on the right depict selected time points of the z-scan time-lapse which was carried out for at least 1 hour following initial photoactivation (for time-lapse, the laser was tuned back to 920 nm with intensity lower than 1%, one z-scan every 3 minutes). Following z-stack image registration, direct comparison of GFP signal between time points was examined manually. Arrowheads point to fusion events, which are recognizable by the abrupt decrease in GFP intensity in photoactivated mitochondria due to GFP dilution into the newly appearing (fusing) mitochondria. Mitochondria that were identified for simply moving away from the photoactivated ROI and did not satisfy these parameters were not considered in our quantification. Bar, 20 µm. **(C)** Quantification of fusion rates in astrocytic end-feet and branches over the course of 1 hour of imaging (n= 17 astrocytes from 4 mice). Note the overall low fusion rate under resting conditions in both branches and end-feet. **(D)** Schematic showing AAV-mediated cytoGCaMP6f expression in astrocytes followed by 2PLSM in slices and subsequent AstroSparks analysis. **(E)** Example of cytoGCaMP6-expressing astrocytes in brain slice following AstroSparks processing and ROI detection (ROIs in end-feet are depicted in white). Right panels depict ROI traces and corresponding raster plots. Bar, 20 µm. **(F)** Quantification of cytosolic Ca^2+^ transients in astrocytic end-feet of wild-type (Ctrl), Mfn2^cKO^ and Mfn1^cKO^ astrocytes under the indicated conditions (n≥ 20 cells from 2-3 different mice per time and condition). **(G)** Frequency of cytosolic and mitochondrial Ca^2+^ transients within branches of Mfn2^cKO^ astrocytes quantified utilizing the indicated sensors and under the specified conditions (n≥ 20 cells from 2-3 mice per time and condition). *, p < 0.05, **, p < 0.01, ***, p < 0.001 (non-parametric Mann-Whitney t-test).

**Figure S6. Related to Figure 6. Assessment of cell proliferation in injured Mfn2^cKO^ mice. (A)** Example of reconstructed cortical vascular network following filament tracing (in white). Systematic inspection of the traced network led to the identification of potential artifacts (false segments, in yellow), which were corrected by manual selection and subsequent elimination. Bar, 100 µm. **(B)** Top views of control and Mfn1^cKO^ cleared cortices showing the extent of dextran-filled vasculature at 7 days post-SW. Arrowheads point to the lesion track. Insets depict zooms of the lesioned core region (circled in white). Bar, 200 µm. **(C)** Quantification of branch points, fractional volume and total length of the vascular network in Mfn1^WT^ and Mfn1^cKO^ cortices (n= 3-4 mice/condition; two-way ANOVA followed by Tukey’s post-hoc test). **(D)** Pictures depicting large views of the injured cortex in Mfn2^WT^ and Mfn2^cKO^ mice at 7 days post-SW and following immunostaining for the nuclear marker SOX2 (labeling astrocytes) as well as EdU. Insets show co-localization of EdU with SOX2 (indicated by yellow arrowheads). Bar, 100 µm. **(E)** Quantification of total proliferating cells (upper graph) as well as proliferating astrocytes (SOX2+/EdU+) within the area surrounding the lesion track in Mfn2^WT^ and Mfn2^cKO^ mice at 7 days post-SW (n= 4-5 mice/condition; nonparametric Mann-Whitney t-test). **(F)** Fraction of endothelial (ERG+) as well as astrocytic (SOX2+) cells being double positive for EdU within the area surrounding the lesion track in Mfn2^WT^ and Mfn2^cKO^ mice at 7 days post-SW (n= 4-5 mice/condition; nonparametric Mann-Whitney t-test). *, p < 0.05, **, p < 0.01, ***, p < 0.001.

**Figure S7. Related to Figure 6 and 7. Analysis of astrocytic mitochondrial morphology and Ca^2+^ dynamics following synthetic linker expression. (A)** Examples of Mfn2^cKO^ brain sections (obtained from Mfn2^lox/lox^ x Glast::CreERT2 x R26^LSL-tdTomato^) and corresponding control samples bearing tdTomato fluorescence in astrocytes. Brain sections were immunostained for the endothelial marker CD31 in order to reveal the vascular network at 7 days following SW-injury. Bar, 100 µm. **(B)** Zoom of reactive tdTomato+ astrocytes at 7 days post-SW in close proximity to the lesion. Pictures depict the polarized morphology of Mfn2^WT^ and Mfn2^cKO^ reactive astrocytes and a comparable extent of perivascular wrapping around the CD31+ vessels. Bar, 10 µm. **(C)** Quantification of astrocyte density within the injured cortices of Mfn2^WT^ and Mfn2^cKO^ tdTomato-expressing mice (n= 4 mice per condition; nonparametric Mann-Whitney t-test). **(D)** Quantification of perivascular mitochondrial fragmentation (i.e., fraction of perivascular fragmented mitochondria as calculated in Figure 2D) in Mfn2^cKO^ astrocytes following transduction with AAV-ctrl or AAV-linker (n= 3 mice per condition; one-way Anova followed by Dunn’s multiple comparison). **(E)** Scheme showing the AAV constructs utilized to express mitoGCaMP6f in tandem with OMM-mRFP-ER (or its control, OMM-mRFP) in astrocytes. **(F)** Experimental design showing AAV delivery followed by 2PLSM in slices and mitochondrial Ca^2+^ uptake analysis via AstroSparks. **(G)** Example of mitoGCaMP6-expressing Mfn2^cKO^ astrocytes in brain slice transduced with either AAV-linker or AAV-ctrl and following AstroSparks processing and ROI analysis (ROIs in the end-feet are depicted in white, while the processes appear in red). Right panels depicts ROI traces and corresponding raster plots. Bar, 20 µm. **(H)** Quantification of the amplitude and duration of mitochondrial Ca^2+^ events in branches and end-feet of Mfn2^cKO^ astrocytes transduced with the AAV-linker or AAV-ctrl (n≥ 29 cells per time and condition). **(I)** Analysis of branching points and total vessel length in CD31 immunostained brain sections obtained from Mfn2^WT^ and Mfn2^cKO^ injured mice (n= 3 mice/condition; nonparametric Mann-Whitney t-test). *, p < 0.05, ns, not significant.

## Methods

### Animal subjects

Six to 8-week old C57BL/6 and transgenic mice of mixed genders were used for stereotactic injections, SW-injury, tamoxifen treatments, slice and *in vivo* imaging. Mice were housed in groups of up to 5 animals per cage supplied with standard pellet food and water *ad libitum* with a 12 h light/dark cycle, while temperature was controlled to 21-22°C. Mice carrying the loxP-flanked genes *Mfn1*^fl/fl^ (Lee et al., 2012) and *Mfn2*^fl/fl^ (Lee et al., 2012) were crossed with the inducible hGFAP-Cre^ERTM^ (Chow et al., 2008) line and subsequently to the Cre-dependent mitochondrial-targeted mitoYFP (Sterky et al., 2011) or GCamp3 reporter (Zariwala et al., 2012). For validation experiments, Mfn2^fl/fl^ mice were crossed with the astrocyte-specific Glast-Cre^ERT2^ line (Mori et al., 2006) in combination with the inducible tdTomato reporter (Madisen et al., 2010). For experiments involving the use of an EnvA-modified Rabies virus to express fluorescent indicators specifically in astrocytes, hGFAP-TVA mice (Holland and Varmus, 1998) expressing the avian membrane-bound TVA receptor under the control of human GFAP promoter were used. All experimental procedures were performed in agreement with the European Union and German guidelines and were approved by the State Government of North Rhine Westphalia.

### Tamoxifen treatments

Mice were intraperitoneally injected with tamoxifen (40 mg/ml dissolved in 90% corn oil and 10% ethanol) once a day for a maximum of 5 consecutive days. All subsequent experiments were performed at least one week after the last tamoxifen injection. The exact time frames are indicated in the text for individual experiments.

### Stereotactic procedures and viral injections

Mice were anesthetized by intraperitoneal injection of a ketamine/xylazine mixture (130 mg/kg body weight ketamine, 10 mg/kg body weight xylazine), treated subcutaneously with Carprofen (5 mg/kg) and fixed in a stereotactic frame provided with a heating pad. A portion of the skull covering the somatosensory cortex (from Bregma: caudal: −2.0; lateral: 1.8) was thinned with a dental drill avoiding to disturb the underlying vasculature. For unilateral SW-injury, a stainless steel lancet was slowly inserted into the cortex to a depth of 0.8 mm, moved 1 mm caudally and then slowly removed. For virus injection a finely pulled glass capillary was inserted through the dura (−0.6 to −0.3 from Bregma) and a total of 200-300 nl of virus were slowly infused via a manual syringe (Narishige) in multiple vertical steps spaced by 50-100 µm each during a time window of 10-20 minutes. After infusion, the capillary was left in place for few additional minutes to allow complete diffusion of the virus. After capillary removal, the scalp was sutured and mice were placed on a warm heating pad until full recovery. Physical conditions of the animals were monitored daily to improve their welfare before euthanize them. For cranial window implantation, anesthetized mice received a pre-emptive subcutaneous injection with Carprofen (5 mg/kg) and dexamethasone (0.25 mg/kg). The scalp was removed and the underlying connective tissue was cleared from the skull. A circular craniotomy (3 mm in diameter) was performed over the posterior parietal cortex using a dental drill and avoiding to disturb the underlying vasculature. During the whole procedure, a saline solution was flushed onto the area exposed with the craniotomy. A sterile 3 mm circular glass coverslip (#1 thickness, Warner Instruments) was gently implanted into the craniotomy site and sealed in place with a thin layer of Sylgard (Sigma) before applying dental cement (Dentalon plus, Heraeus Kulzer GmbH) to fix the coverslip and cover the surrounding exposed skull. An aluminium chamber plate (CP-1, Narishige) was fixed with cement on top of the cover glass to facilitate mouse head immobilization at the 2-photon microscope via a head holder (MAG-2, Narishige). A single tail vein injection of 50 µl Dextran Texas Red (70 kDa, Thermo Fisher, D1864) in saline was used to label the brain vasculature in anesthetized animals. The depth of anaesthesia was assessed throughout the surgery and recording time (usually 1-2 hours) and eventually mice received one or more additional boluses of anaesthetic each corresponding to one third of the initial dose.

### Viral production

Construction of the glycoprotein (G protein) gene-deleted RABV (SADΔG-mCherry) and virus rescue from pHH-SADΔG-mCherry SC has been described before (Ghanem et al., 2012). cDNAs encoding organelle-targeted fluorescent protein genes were used replace the mCherry ORF of using unique NheI/NotI restriction sites. RABVΔG-mito-tagRFP and RABVdG-mito-PA GFP contains the pre-peptide of human ornithine carbamoyltransferase fused to the N terminus of tagRFP (Yi et al., 2017) or PA-GFP. ER-targeted GFP contains an N-terminal ER retention sequence (KDEL-GFP, kindly provided by E. Snapp). Viruses pseudotyped with the homologous SAD G glycoprotein were amplified in BSR MG-on cells complementing the G deficiency of the virus upon induction of G expression by doxycycline (Finke et al., 2003) and viruses pseudotyped with the EnvA protein in BHK-EnvARGCD cells expressing an ASLV-A envelope protein comprising the RABV G cytoplasmic tail (Wickersham et al., 2007). The G- or EnvA-coated virus was concentrated by ultracentrifugation and used for in vivo injection. Plaque-forming unit (pfu) number titration was performed by infecting BHK-wt cells and HEK293T-TVA cells with G-coated virus and EnvA-coated virus, respectively. Helper-free AAV vectors were either obtained from Vector Biolabs as custom projects or produced according to standard manufacturer’s instructions (Cell Biolabs). Briefly, 293AAV cells were transiently transfected with a transfer plasmid carrying the desired transgenes along with a packing plasmid encoding the AAV1 capsid proteins and a helper plasmid, using the calcium phosphate method. Crude viral supernatants were obtained via lysing cells in PBS by freeze-thaw cycles in a dry ice/ethanol bath. The AAV vectors were purified by discontinuous iodixanol gradient ultracentrifugation (24h at 32,000 rpm and 4°C) and concentrated using Amicon ultra-15 centrifugal filter unites. Genomic titres were determined by real-time qPCR.

### In vivo and ex vivo imaging

Isolated brains were placed in ice-cold, carbogen-saturated (5% CO_2_, 95% O_2_, pH 7.4) artificial cerebrospinal fluid (ACSF) containing (in mM): 125 NaCl, 2.5 KCl, 1.25 NaH_2_PO_4_, 25 NaHCO_3_, 25 Glucose, 0.5 CaCl_2_ and 3.5 MgCl_2_ (osmolarity of 310-330). 270-300 µm thick coronal slices were obtained using a vibratome (Micron, HM 650V) and transferred into a pre-incubation chamber maintained at room temperature and containing ACSF supplemented with 1 mM CaCl_2_ and 2 mM MgCl_2_. During imaging, slices were moved in a dedicated imaging chamber and experiments were conducted under continuous ACSF perfusion at a constant temperature of 32-33°C. Imaging in slices and *in vivo* was performed using a multiphoton laser-scanning microscope (TCS SP8 MP-OPO, Leica Microsystems) equipped with a Leica 25x objective (NA 0.95, water) and a Ti:Sapphire laser (Chameleon Vision II, Coherent). For Calcium imaging, detection of fluorescence changes of the GCaMP6f sensor in single astrocytes was achieved by tuning the laser to 920 nm. This wavelength also allowed simultaneous recording of Dextran Red signal in experiments *in vivo*. Two internal HyD detectors (FITC: 500-550 nm, TRITC: 565-605 nm) were utilized to monitor GCaMP6 and Dextran Red signals. Typical recording sessions consisted in 3-5 min of continuous imaging (resolution of 1024×1024 pixels and zoom of 1 or 5) with a frame rate of 1.16 frames /s (0.86 s/frame). Analysis of Ca^2+^ transients acquired with higher frame rates (up to 10 Hz) yielded comparable results in terms of frequency, amplitude and duration of events, but worsened the overall image quality. For mito-PA-GFP experiments, photoactivation of selected ROIs of individual astrocytes was carried out by tuning the 2-photon laser to 840 nm (10% of laser power for 10-20 seconds), while time-lapse imaging was performed utilizing GFP excitation (920nm) and an internal HyD detector (FITC: 500-550 nm). Usually 2-3 ROIs of identical size per astrocytes were selected in the end-feet and branches and, after photoactivation, the whole astrocyte volume (inter-stack interval of 1 μm) was imaged over the course of at least 1h every 3 minutes. Only astrocytes located at least 20-30 μm below the slice surface, with a general healthy appearance throughout the recording time (i.e., absence of visibly fragmented mitochondria) and whose acquisitions displayed only no or a minor spatial drift in xyz during the whole imaging session were included in subsequent analysis. Acquired time points were then merged in a 4D hyperstack in ImageJ and the resulting 3D volumes registered utilizing the “Correct 3D drift” plugin in ImageJ. Quantification of fusion events was performed manually by inspecting the volumes including and surrounding the photoactivated ROIs. Fusion events were identified by the abrupt decrease in GFP intensity in directly photoactivated mitochondria due to GFP dilution into the newly appearing (fusing) mitochondria that were not initially photoactivated. In rare cases, mitochondria that simply moved away or though the photoactivated ROIs and did not satisfy these fusion parameters were not considered in our quantification.

### Calcium imaging analysis via AstroSparks

Time-lapse image sequences were drift-corrected by utilizing the “fast & rigid body” options of the TurboReg plugin (http://bigwww.epfl.ch/thevenaz/turboreg/) in ImageJ, aligning each frame to a median projection of eleven frames centered on the middle of the time series. In case of non-satisfactory results, the moco plugin (https://github.com/NTCColumbia/moco) was used alternatively. The image sequence was then cropped to exclude border regions that were not acquired throughout the whole recording period. The noise was reduced with an isotropic (σ = 2 px, xyt) Gaussian Blur filter. Next, only pixels with a median intensity or a peak intensity in a median filtered (radius: 5 px) and background corrected (Subtract Background plugin, options: “rolling=500 sliding disable”) image exceeding the threshold of 5 (a.u.) were considered for further analysis. Based on a standard deviation (SD) projection, the FindFoci plugin identified regions of interest (ROIs). The threshold was set to the mean + 3x SD intensity of pixels identified as background by the “IsoData” auto-threshold. ROIs included all neighboring pixels with an intensity higher than a per ROI threshold of: (maximum intensity – background) x 0.4 + background, to compensate for the spreading of bright signals. ROIs smaller than 0.3 µm^2^ were excluded (in case of the mitoGCaMP6 script, the plugin was used on a median projection to include high, yet stable signals, meaning all mitochondrial ROIs). Next the ΔF/F was calculated based on a median projection reference. ROIs with a high ΔF/F were additionally identified by the FindFoci plugin on a mean filter (radius: 5 px) smoothed maximum projection. Finally, all ROIs were projected onto each other and overlapping ROIs were combined. Once ROIs were identified, the area and average intensity per ROI and time point were handed to IgorPro (v7.0.4.1, WaveMetrics, Inc., Lake Oswego, Oregon 97035, USA). Custom written routines identified the duration, amplitude, and frequency of events deviating from baseline. In order to correct for bleaching, all traces were averaged and fitted with an exponential decay function. Based on this reference all traces were corrected. The baseline was identified as follows: the average intensity per ROI was smoothed with a mean-sliding box algorithm (width: 3 time frames). The obtained values were sorted in ascending order and for each rank the standard deviation including all lower ranking values was calculated. To define the threshold at which the SD suddenly increases, i.e. when values start to deviate from baseline and thus increase the SD, the difference in SD (smoothed with a mean-sliding box algorithm (width: 3 frames)) between subsequent ranks was calculated. The rank, at which half maximal difference was reached for the first time, marks the threshold. If the threshold contained less than 15% of all values, the whole trace was defined as baseline. Its mean was used to calculate ΔF/F. In order to identify events, the 20% quantile (of 7 sliding frames) needed to exceed the 80% quantile (of 11 sliding frames) with a time lag of 2.58 s (3 frames) by 1.5 x the SD of the low-cut frequency-filtered (0.2 Hz) ΔF/F trace. For all those events, the end was defined as the earlier time point at which the ΔF/F trace crossed zero or crossed the ΔF/F level just prior to the start of the event.

### Tissue clearing

To assess the structure of the organelles and vasculature in intact cortices, the tissue was cleared using the short Sca*l*eS protocol described previously (Hama et al., 2011). Following brain isolation and overnight post-fixation in 4% PFA at 4°C, the ventral portion of the brain was removed and the remaining dorsal part (including the somatosensory cortex) was placed in Sca*l*eSQ(5) solution for 1 d at 37°C followed by incubation in Sca*l*eS4(0) for another day at 37°C. Sca*l*eSQ(5) was composed of 22.5% D-(-)-sorbitol (w/v), 9.1 M Urea, 5% Triton X-100 (w/v), pH 8.2 and Sca*l*eS4(0) of 40% D-(-)-sorbitol (w/v), 10% Glycerol (w/v), 4 M Urea, 15-25% DMSO (v/v), pH 8.1. The next day, cleared cortices were placed in an imaging chamber filled with Sca*l*eS4(0).

### 3D Reconstructions and analysis

For analysis of ER-GFP-expressing astrocytes and vascular networks in 3D, 2PLSM (TCS SP8 MP-OPO, Leica Microsystems, 25x water immersion Objective, resolution of 1024×1024 pixels, zoom factor of 2, frame average of 2 and 1 µm inter-stack interval) was utilized to acquire the desired volumes in cleared cortices and the resulting z-stacks were imported into the Imaris software (version 8.3.1, Bitplane) to obtain a rendered 3D volume utilizing the acquisition parameters. A surface mask was generated from the resulting volumes of the desired fluorescent signal. To analyse the fractional ER-GFP signal density, the created mask was then utilized to carefully segment the different cellular compartments, including the soma, end-feet (corresponding to the portion of ER-GFP signal surrounding dextran-labelled vessels) and by exclusion the branches. ER-GFP integrated signal density was then calculated for each of these sub-volumes per cell and condition. For analysis of vasculature density in cleared control and injured cortices as shown in Figure 6, z-stacks were acquired at a resolution of 1024×1024 pixels (whole fields of view, 590 x 590 μm scanning from surface of the cortex to the beginning of white matter) with a zoom factor 0.75 and 2 µm inter-stack interval. Magnifications were acquired with a zoom factor of 2, a frame average of 2 and a 1µm inter-stack interval. Large volume acquisitions imported into Imaris were first cropped in xy on both sides of the lesion track in order to obtain a narrower area of 590 x 350 μm (175 μm on each side of the lesion along the whole track utilizing the dextran signal as reference), and then cropped in z to 600 μm to obtain a final cortical block of 590 x 350 x 600 μm. A surface mask was then generated from the dextran fluorescent signal and utilized to trace the vascular network via a filament tracing algorithm embedded in Imaris. Following filament tracing, volumes were thoroughly inspected for potential artifacts and eventually corrected (see Figure S6A) before extracting vascular fractional volume, total length and branching points. For quantifications of vascular network complexity following expression of AAVs as shown in in Figure 7H and S7I, the AngioTool ImageJ plugin (Zudaire et al., 2011) was used.

### Immunostainings

Following overnight post-fixation of isolated brains with PFA 4% in PBS, coronal brain sections (40 to 70 µm thick) were prepared using a vibratome (Leica, VT1000 S) and permeabilized in 1% Triton X-100 in PBS for 10 min at RT, followed by brief incubation in 5% BSA and 0.3% Triton X-100 in PBS before overnight immunodetection with primary antibodies diluted in blocking buffer at 4°C on an orbital shaker. The next day, sections were rinsed in PBS 3x 10 min and incubated for 2h at RT with the respective fluorophore-conjugated secondary antibodies diluted in 3% BSA. After washing and nuclear counterstaining with 4’,6-diamidino-2-phenylindole (DAPI, ThermoFisher, 3 µM), sections were mounted on microscopic slides using Aqua Poly/Mount (Polysciences). The following primary antibodies were used: chicken anti-GFP (1:500, Aves Labs, GFP-1020), rabbit anti-RFP (1:500, Rockland, #600401379), rabbit anti-GFAP (1:500, Millipore, ab5804), mouse anti-GFAP (1:500, Millipore, MAB360), rat anti-CD45 (1:500, BD, #550539), rat anti-CD31 (1:50, BD, #550274), rabbit anti-S100β (1:500, Millipore, ab52642). The following secondary antibodies were used (raised in donkey): Alexa Fluor 488-, Alexa Fluor 546-, Alexa Fluor 647-conjugated secondary antibodies to rabbit, mouse, chicken and rat (1:1000, Jackson ImmunoResearch). Images were acquired utilizing a SP8 Confocal microscope (Leica) equipped with a 20x (NA 0.75), 40x (NA 1.3), 63x (NA 1.4) or 100x (NA 1.3) oil immersion objective and further processed with Fiji.

### Isolation and enrichment of astrocytes via magnetic cell sorting (MACS)

For astrocyte enrichment the kit “Isolation and cultivation of astrocytes from adult mouse brain” (Miltenyi Biotec) was used according to the manual instructions. In brief, cortical brain tissue was extracted and dissociated enzymatically as well as mechanically. Myelin and cell debris were eliminated and in a subsequent step erythrocytes were removed. Using the anti-ACSA-2 microbeads with the autoMACS Pro Separator, astrocytes were magnetically separated from the suspension. Enriched astrocytes were further processed by mass spectrometry.

### Isolation and enrichment of astrocytes via fluorescence-activated cell sorting (FACS)

The cortical region exposed to SW-injury (typically 1 mm wide, 2 mm long and spanning the cortex depth but excluding the white matter) was microdissected and dissociated using the “Adult Brain Dissociation” kit from Miltenyi Biotec following the manufacturer’s instructions. Following astrocyte staining with ACSA-2-APC (Clone IH3-18A3, 1:200, Miltenyi Biotec), Hoechst to discriminate between cells and debris (10 µg/ml, Cell Signaling) and 7AAD for viability (2.5 µg, Affymetrix), 20.000 events of control samples were recorded to set appropriate gates and compensations. Cell sorting was performed with a BD FACSAria Fusion equipped with a 100 µm nozzle (20 psi) and five lasers (UV 355 nm, violet 405 nm, blue 488 nm, yellow 561 nm and red 640 nm). Sorted cells were collected in PBS and processed for mass spectrometry analysis.

### Transmission electron microscopy and image analysis

Anesthetized mice were transcardially perfused with a fixative solution containing 4% formaldehyde and 2.5% glutaraldehyde in 0.1 M cacodylate buffer. The brain was isolated, cut in 1 mm thick sagittal sections and small portions of upper layers of the cortex were dissected for further processing (for injured cortices, the examined area was dissected according to the location of the lesion track). For EPON embedding, the fixed tissue was washed with 0.1 M sodium cacodylate buffer, incubated with 2% OsO_4_ in 0.1 M cacodylate buffer (Osmium, Science Services; Caco Applichem) for 2 h at 4°C and washed again three times with 0.1 M cacodylate buffer. Subsequently, tissue was dehydrated using an ascending ethanol series with 15 min incubation at 4°C in each EtOH solution. Tissues were transferred to propylene oxide and incubated in EPON (Sigma-Aldrich) overnight at 4°C. Tissues were placed in fresh EPON at RT for 2 h, followed by embedding for 72 h at 62°C. Ultrathin sections of 70 nm were cut using an ultramicrotome (Leica Microsystems, UC6) with a diamond knife (Diatome, Biel, Switzerland) and stained with 1.5% uranyl acetate at 37°C for 15 min and lead citrate solution for 4 min. For immunogold staining (Tokuyasu technique), fixed tissue (4% PFA and 0.2% GA) was infiltrated with 2.3 M sucrose in 0.1 M phosphate buffer overnight at 4°C, mounted on aluminium pins for cryo-ultramicrotomy and snap-frozen in liquid nitrogen. Ultrathin cryo-sections of 70 nm were cut with a diamond knife (Diatome, Biel, Switzerland) using a Leica UC6 with FC7 at −90°C. Sections were picked up in a 1:1 mixture of 2% methylcellulose (Sigma-Aldrich) and 2.3 M sucrose. After rinsing 3x in PBS and incubation in 0.05 M glycine (Sigma-Aldrich), sections were blocked (2×3 min) with 1% BSA in PBS. For immuno-labelling, sections were incubated with antiserum specific for RFP (Rockland) in blocking buffer, followed by rinsing of 6x in PBS and 90 min incubation with protein A-gold (12 nm, CMC-Utrecht) in blocking buffer. After fixation with 2% glutaraldehyde (3 min), sections were washed in PBS and H_2_O and contrasted (5 min) with uranyl acetate (0.4% in 2% methylcellulose) on ice. Sections were picked up with a wire loop. Excess fluid was drained from the loop by gentle tapping to Whatman filter paper, and sections were embedded in the remaining thin film by air-drying. Electron micrographs were taken with a JEM-2100 Plus Transmission Electron Microscope (JEOL), equipped with Camera OneView 4 K 16 bit (Gatan) and software DigitalMicrograph (Gatan). For analysis, electron micrographs were acquired with a digital zoom of 5000x or 6000x. The area, perimeter and circularity of each mitochondrion was determined in ImageJ follow manual drawing of single organelles. Mitochondrial density was assessed by quantifying the absolute number of mitochondria per measured astrocytic area or length of basal lamina. A similar approach was utilized to quantify the extent of mitochondria-ER contact sites (defined as sites of contact within a reciprocal distance of 50 nm) and minimal mitochondria-ER proximity. All parameters obtained from one field of view (usually containing several mitochondria and multiple contact sites) were averaged together.

### Mass spectrometry (MS) and data analysis

For proteomic analysis, MACS-enriched or FACS isolated astrocytes were lysed in SP3 lysis buffer (4% SDS in PBS) and chromatin was degraded using a Bioruptor (10 min, cycle 30/30 s). Samples were reduced with 5 mM Dithiothreitol (DTT) at 55°C for 30 min, alkylated with 40 mM Chloroacetamide (CAA) at RT for 30 min and protein amount was quantified using the Direct Detect Spectrometer from Merck. Protein digestion was performed using the Single-Pot Solid-Phase-enhanced Sample Preparation approach SP3. In brief, 2 µL of a 10 mg/mL mixture of hydrophilic and hydrophobic carboxylate coated paramagnetic beads (SeraMag Speed Beads, #44152105050250 and #24152105050250, GE Healthcare) were added to each sample. Acetonitrile was added to a final concentration of 50%. Bound proteins were washed with 70% ethanol and 100% acetonitrile. Beads were re-suspended in 5 µL 50 mM Triethylammoniumbicarbonate buffer containing 0.1 µg Trypsin (Sigma) and 0.1 µg LysC (Wako). Digestion was carried out at 37°C for 16 h in a PCR cycler. Recovered peptides were re-suspended in 1% formic acid / 5% DMSO and stored at −20°C prior MS analysis. All samples were analyzed on a Q-Exactive Plus (Thermo Scientific) mass spectrometer that was coupled to an EASY nLC 1000 UPLC (Thermo Scientific). Peptides were loaded with solvent A (0.1% formic acid in water) onto an in-house packed analytical column (50 cm × 75 µm I.D., filled with 2.7 µm Poroshell EC120 C18, Agilent). Peptides were chromatographically separated at a constant flow rate of 250 nL/min using the following gradient: 5-30% solvent B (0.1% formic acid in 80% acetonitrile) within 65 min, 30-50% solvent B within 13 min, followed by washing and column equilibration. The mass spectrometer was operated in data-dependent acquisition mode. The MS1 survey scan was acquired from 300-1750 m/z at a resolution of 70,000. The top 10 most abundant peptides were isolated within a 2 Da window and subjected to HCD fragmentation at a normalized collision energy of 27%. The AGC target was set to 5e5 charges, allowing a maximum injection time of 110 ms. Product ions were detected in the Orbitrap at a resolution of 17,500. Precursors were dynamically excluded for 20 s. All mass spectrometric raw data were processed with Maxquant (version 1.5.3.8) using default parameters. Briefly, MS2 spectra were searched against the Uniprot MOUSE.fasta database, including a list of common contaminants. False discovery rates on protein and PSM level were estimated by the target-decoy approach to 0.01% (Protein FDR) and 0.01% (PSM FDR), respectively. The minimal peptide length was set to 7 amino acids and carbamidomethyolation at cysteine residues was considered as a fixed modification. Oxidation (M) and Acetyl (Protein N-term) were included as variable modifications. The match-between runs option was enabled. LFQ quantification was enabled using default settings. The Maxquant output was processed as follows: Protein groups flagged as „reverse“, “potential contaminant” or “only identified by site” were removed from the proteinGroups.txt. LFQ values were log2 transformed. Proteins with less than 2 valid values were removed. Missing values were replaced by imputation from a normal distribution (width 0.3, down shift 1.8). A two sample t-test was used to determine significantly changing protein levels (S0 = 0.1), and a permutation-based FDR was calculated to correct for multiple testing. The obtained data was uploaded into the Ingenuity Pathway Analysis (IPA) software (Qiagen) utilizing a Benjamini adjusted *p*-value of 0.05 or lower to investigate canonical pathways that were significantly changed. Heat map visualization of relative protein abundance was obtained calculating a z-score of the LFQ values for each protein.

### ^13^C-glucose feeding in mice and sample preparation

Mice were fasted overnight before being anesthetized and receiving a single tail vein injection of 150 μmol of ^13^C_6_-glucose (in saline) over the course of 30 seconds. After 30 min, mice were quickly sacrificed and the peri-lesioned cortical area extracted in PBS for astrocyte enrichment via MACS as described above. The resulting astrocytic fraction was homogenized in acetonitrile:methanol:water (40:40:20) for metabolite extraction.

### LC-MS analysis of isotope-enrichments in amino acids after ^13^C-glucose feeding

For amino acid analysis the benzoylchlorid derivatization method (Wong et al., 2016) was used. In brief: One of the two dried metabolite pellets of each sample was re-suspended in 20 µl of the LC-MS-grade waters (Milli-Q 7000 equipped with an LC-Pak and a Millipak filter, Millipore). The re-suspended sample was mixed with 10 µl of 100 mM sodium carbonate (Sigma) followed by the addition of 10 µl 2% benzoylchloride (Sigma) in acetonitrile (Optima-Grade, Fisher-Scientific). Samples were vortexed before centrifuging them for 10 min 21.300x g at 20°C. Clear supernatants were transferred to fresh auto sampler tubes with conical glass inserts (Chromatographie Zubehoer Trott) and analyzed using an Acquity iClass UPLC (Waters) connected to a Q-Exactive HF (Thermo). For the analysis, 2 µl of the derivatized sample were injected onto a 100 x 1.0 mm HSS T3 UPLC column (Waters). The flow rate was set to 100 µL/min using a buffer system consisted of buffer A (10 mM ammonium formate (Sigma), 0.15% formic acid (Sigma) in Milli-Q water (Millipore)) and buffer B (acetonitrile, Optima-grade, Fisher-Scientific). The LC gradient was: 0% B at 0 min; 0-15% B 0-0.1 min; 15-17% B 0.1-0.5 min; 17-55% B 0.5-14 min, 55-70% B 14-14.5 min; 70-100% B 14.5-18 min; 100% B 18-19 min; 100-0% B 19-19.1 min, 19.1-28 min 0% B. The mass spectrometer was operating in positive ionization mode monitoring the mass range m/z 50-750. The heated ESI source settings of the mass spectrometer were: Spray voltage 3.5kV, capillary temperature 250°C, sheath gas flow 60 AU and aux gas flow 20 AU at a temperature of 250°C. The S-lens was set to a value of 60 AU. Data analysis of isotope ratios was performed using the TraceFinder software (Version 4.2, Thermo Fisher Scientific). Identity of each compound was validated by authentic reference compounds, which were analysed independently. For the isotope enrichment analysis the area of the extracted ion chromatogram (XIC) of each isotope [M + H]^+^ were determined with a mass accuracy (<5 ppm) before calculating the proportions of each detected isotope towards the sum of all isotopes of the corresponding compound. These proportions are given as percent values for each isotope. Results are indicated as Molar Percent Enrichment (M.P.E.), which value is obtained by using the formula:

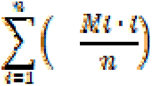

where n=number of carbon atoms in the metabolite, and Mi =relative abundance of the i-th mass isotopomer.

### GC-MS analysis of isotope-enrichments in metabolites from TCA cycle after ^13^C-glucose feeding

Similar to the analysis of the isotope enrichment analysis in the amino acids, isotope enrichment analysis in TCA cycle metabolites were determined using GC-MS (Q-Exactive GC-Orbitrap, Thermo Fisher Scientific). For this purpose metabolites were derivatized using a two-step procedure starting with an methoxyamination (methoxyamine hydrochlorid, Sigma) followed by a trimethyl-silylation using N-Methyl-N-trimethylsilyl-trifluoracetamid (MSTFA, Macherey-Nagel). Dried samples were re-suspended in 5 µL of a freshly prepared (20 mg/mL) solution of methoxyamine in pyridine (Sigma) to perform the methoxyamination. These samples were then incubated for 90 min at 40°C on an orbital shaker (VWR) at 1500 rpm. In the second step additional 45 µL of MSTFA were added and the samples were incubated for additional 30 min at 40°C and 1500 rpm. At the end of the derivatisation the samples were centrifuged for 10 min at 21100x g and 40 µL of the clear supernatant was transferred to fresh auto sampler vials with conical glass inserts (Chromatographie Zubehoer Trott). For the GC-MS analysis 1 µL of each sample was injected using a PAL autosamplee system (Thermo Fisher Scientifc) using a Split/Splitless (SSL) injector at 300 °C in splitless mode. The carrier gas flow (helium) was set to 2 ml/min using a 30m DB-35MS capillary column (0.250 mm diameter and 0.25 µm film thickness, Agilent). The GC temperature program was: 2 min at 85°C, followed by a 15!C per min ramp to 330°C. At the end of the gradient the temperature is held for additional 6 min at 330°C. The transfer line and source temperature are both set to 280°C. The filament, which was operating at 70 V, was switched on 2 min after the sample was injected. During the whole gradient period the MS was operated in full scan mode covering a m/z range between 70 and 800 with a scan speed of 20 Hertz. For data analysis peak areas of extracted ion chromatograms of each isotope of compound-specific fragments [M - e^-^]^+^ were determined using the TraceFinder software (Version 4.2, Thermo Fisher Scientific) with a mass accuracy (<5 ppm). Subsequently proportions of each detected isotope towards the sum of all isotopes of the corresponding compound-specific fragment were determined. These proportions are given as percent values for each isotope. Details on the compound-specific fragments of the analysed compounds: citric acid was analysed from a five carbon-containing fragment (C11H21O4Si2) and a m/z of 273.09729; succinic acid was analysed from a four carbon-containing fragment (C9H19O4Si2) and a m/z of 247.08164; fumaric acid was analysed from a four carbon-containing fragment (C9H17O4Si2) and a m/z of 247.08164. The retention time and therefore identity of each compound was validated by authentic reference compounds which were analysed independently.

### Analysis of mitochondrial morphology, 2D ER morphology and CD31 immunoreactivity

For analysis of mitochondrial morphology in mitoYFP+ samples, serial z-stacks (0.3 to 0.5 µm steps) of individual astrocytes within the slice were acquired with an SP8 laser scanning confocal system (Leica) utilizing a 100x objective (NA 1.3) and digital zoom of 1.5. Acquired z-stacks were subjected to deconvolution (Huygens Professional software; Scientific Volume Imaging) utilizing the acquisition parameters and the resulting surface rendering images were utilized to extract mitochondrial morphological parameters (length, voxel volume and sphericity) via the object analyser plugin (Huygens). The length and sphericity of all quantified mitochondria (typically in the range of several hundreds) per astrocyte were plotted via the OriginPro software (OriginLab) and the resulting diagrams utilized to quantify the percentage of fragmented vs tubular mitochondria per astrocyte, utilizing as cut-off values 1μm for the length and 0.8 for sphericity (where 1 would represent a sphere). At least 4-5 astrocytes (selected for their proximity to the lesion track) per mouse were analysed and the percentage of all individual astrocytes from the same mouse were pooled together. To estimate the perivascular density of mitochondria in astrocytic end-feet, a circular ROI exceeding the dextran-red signal by 5 μm (for cleared tissue) or 2 μm (for brain sections stained with CD31) was drawn around the labelled vessels in each analysed image. The acquired channel containing the mitochondrial signal (mitoYFP or mRFP depending on the experimental setup) was first thresholded and the resulting image utilized to calculate the ROI area fraction covered by mitochondrial signal. To calculate the perivascular ER-GFP *g*-ratio, the thickest sheet of ER-GFP signal in the end-feet in direct contact with the dextran-labelled vessel was measured and normalized to the vessel radius itself. The formula (R_lumen_/(R_lumen_ + R_ER-GFP_)) was utilized to obtain the *g*-ratio values per astrocyte. To analyse the vasculature in 2D in sections labelled for CD31, a region of about 600×600 µm in xy was cropped in the acquired z-stacks, its brightness adjusted with the same parameters for all images, smoothed in 3D (sigma =1) and signal noise removed via a despeckle filter. Following z-projection (standard deviation, STD) and creation of a binary mask, the CD31 area fraction was measured for each image.

### Statistics

Data are represented as means ± SD. Graphical illustrations and significance were obtained with GraphPad Prism 7 (GraphPad) or with OriginPro (OriginLab). The levels of significance were set as * p < 0.05; ** p < 0.01; *** p < 0.001.

### Materials availability

Requests for materials, reagents and tools should be addressed to the lead contact, Matteo Bergami.

