## Supplementary figures and images for "Mitochondria-ER contacts in reactive astrocytes coordinate local perivascular domains to promote vascular remodelling"

### Supplemental Figures 1-7

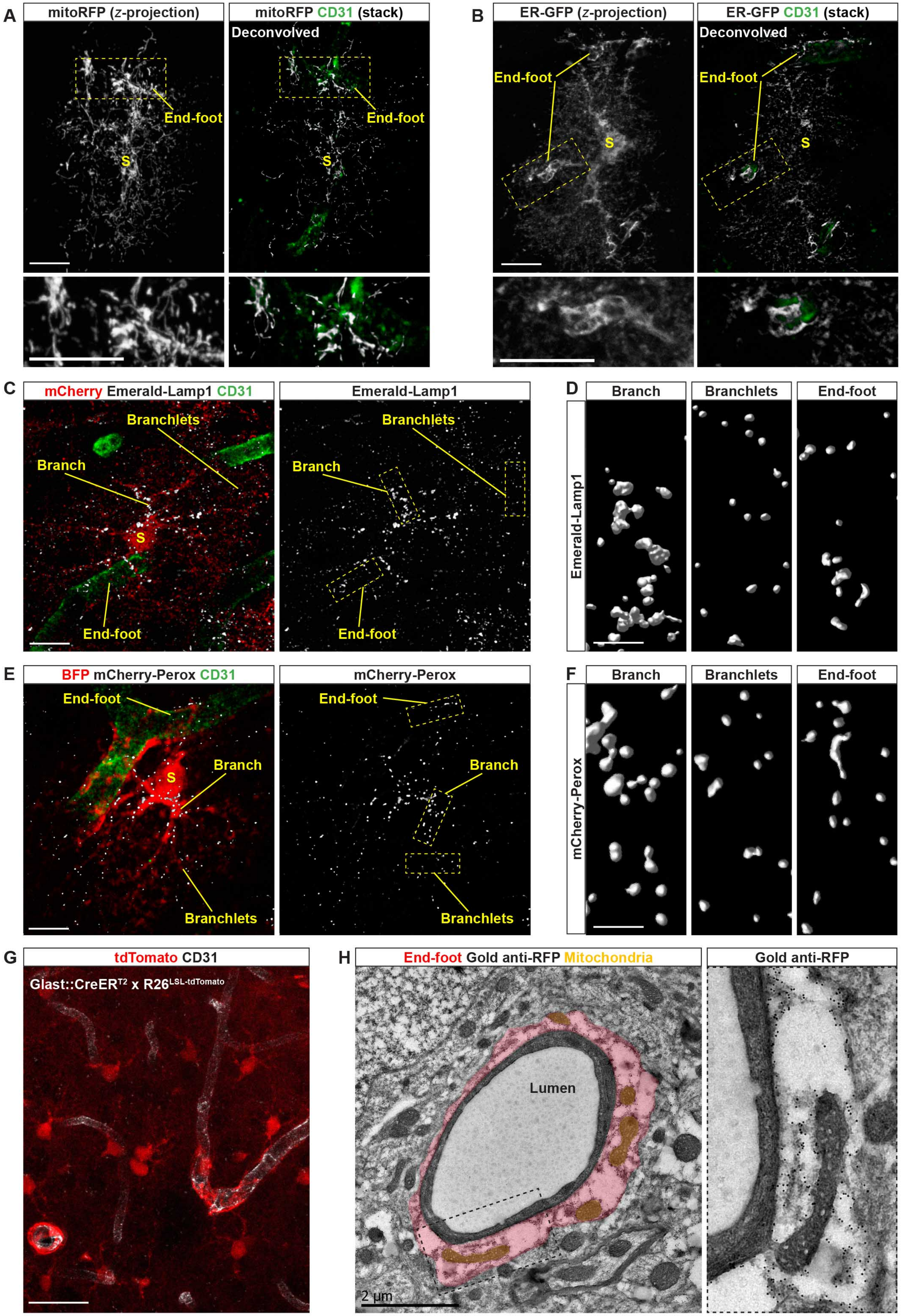

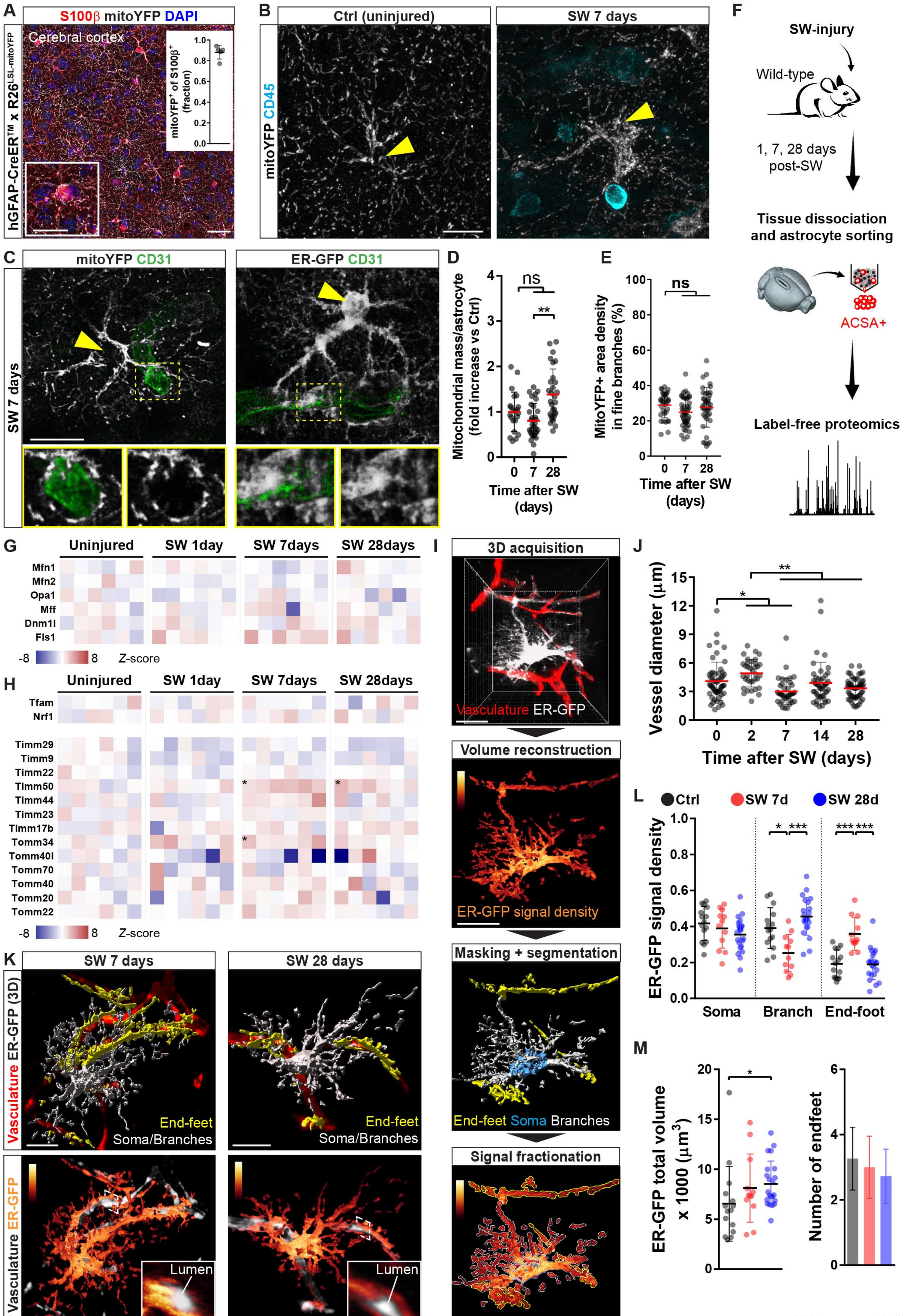

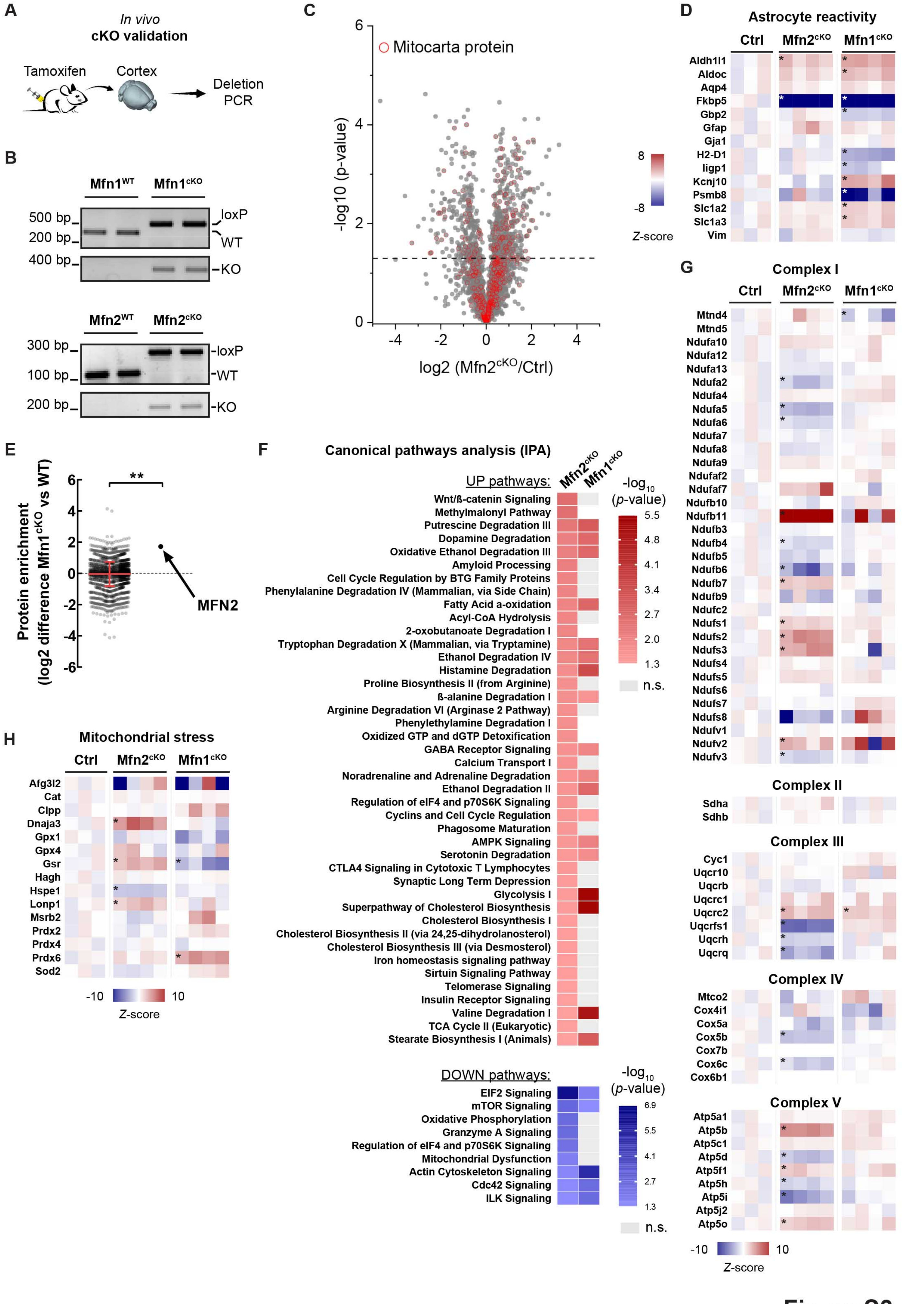

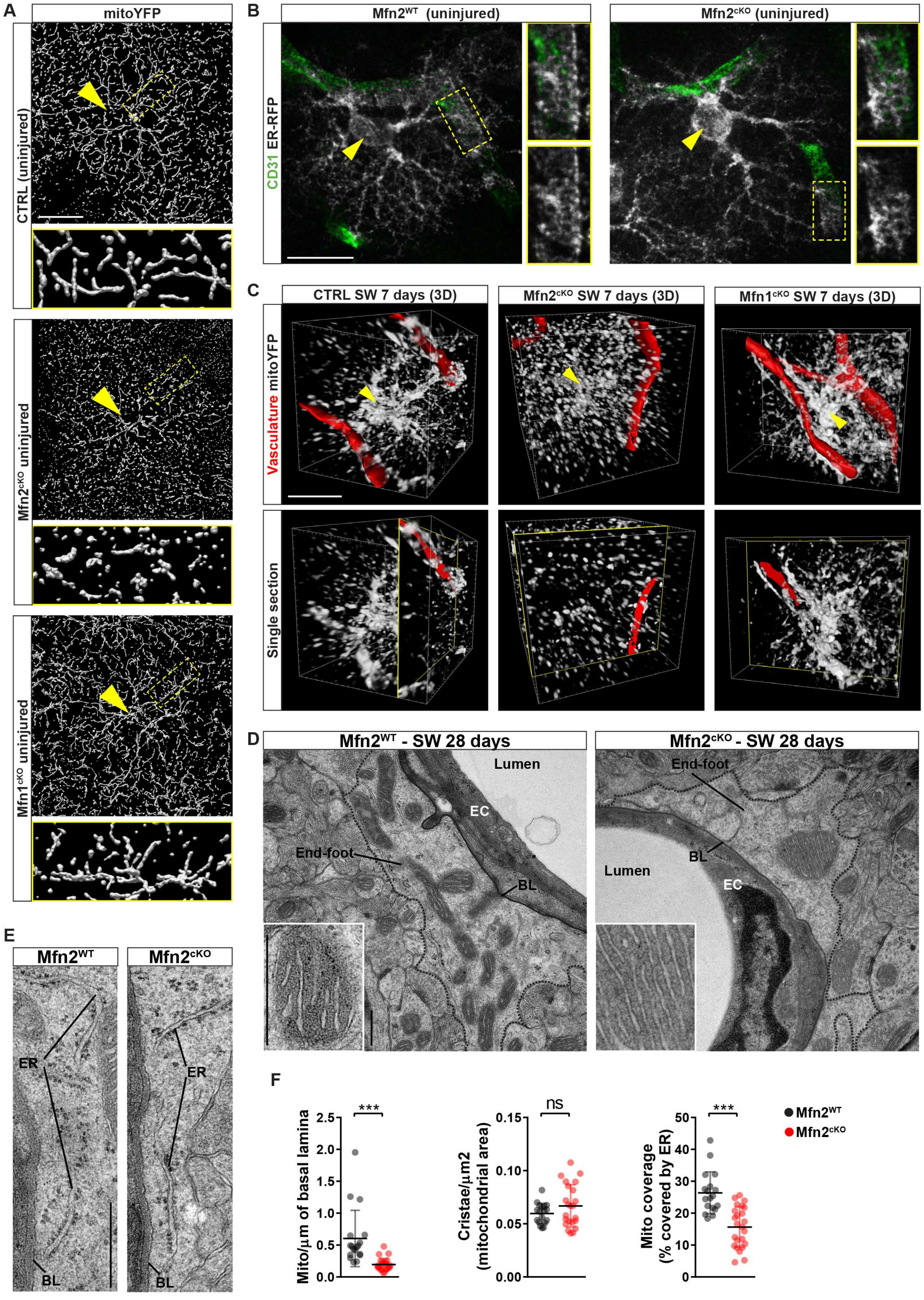

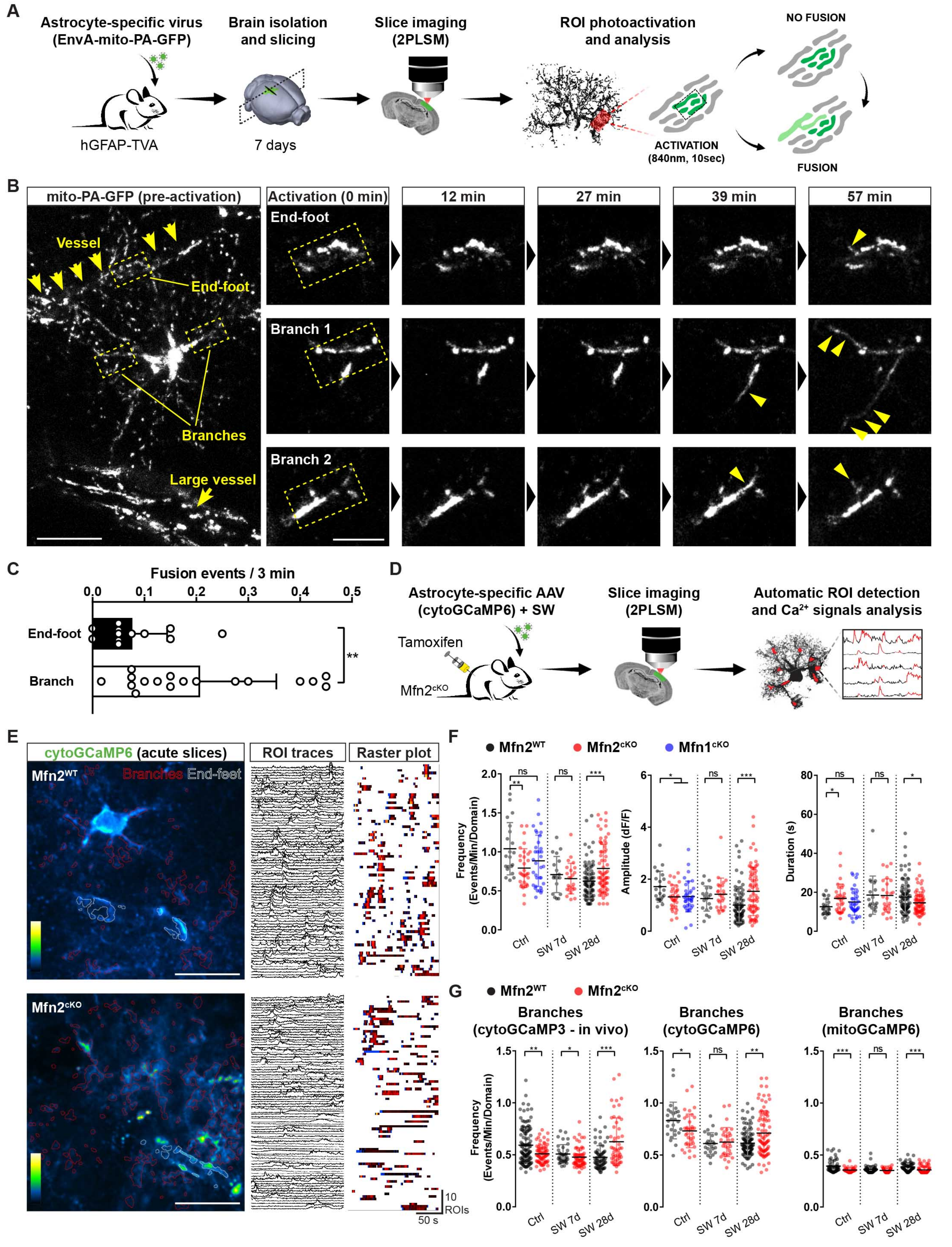

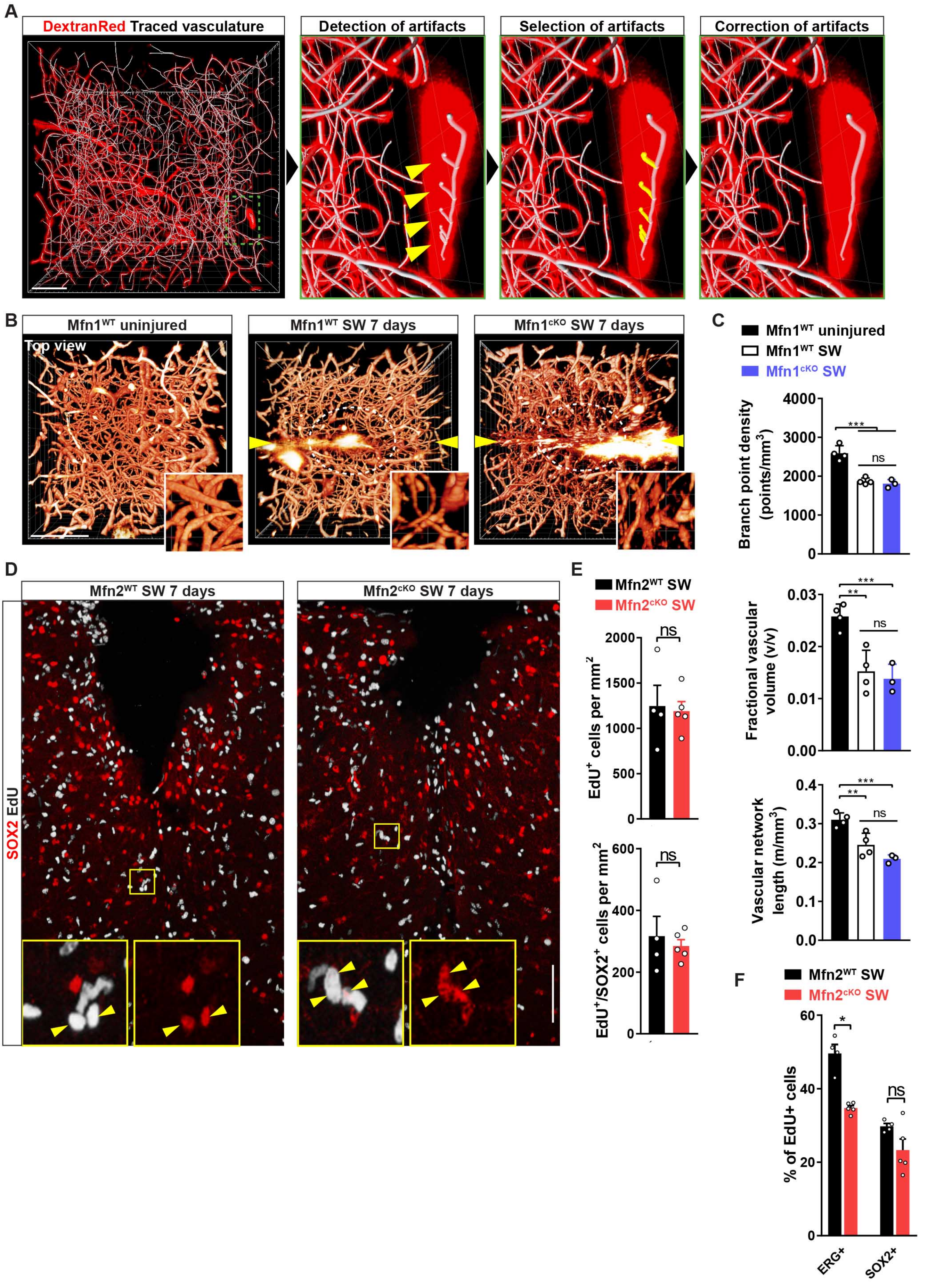

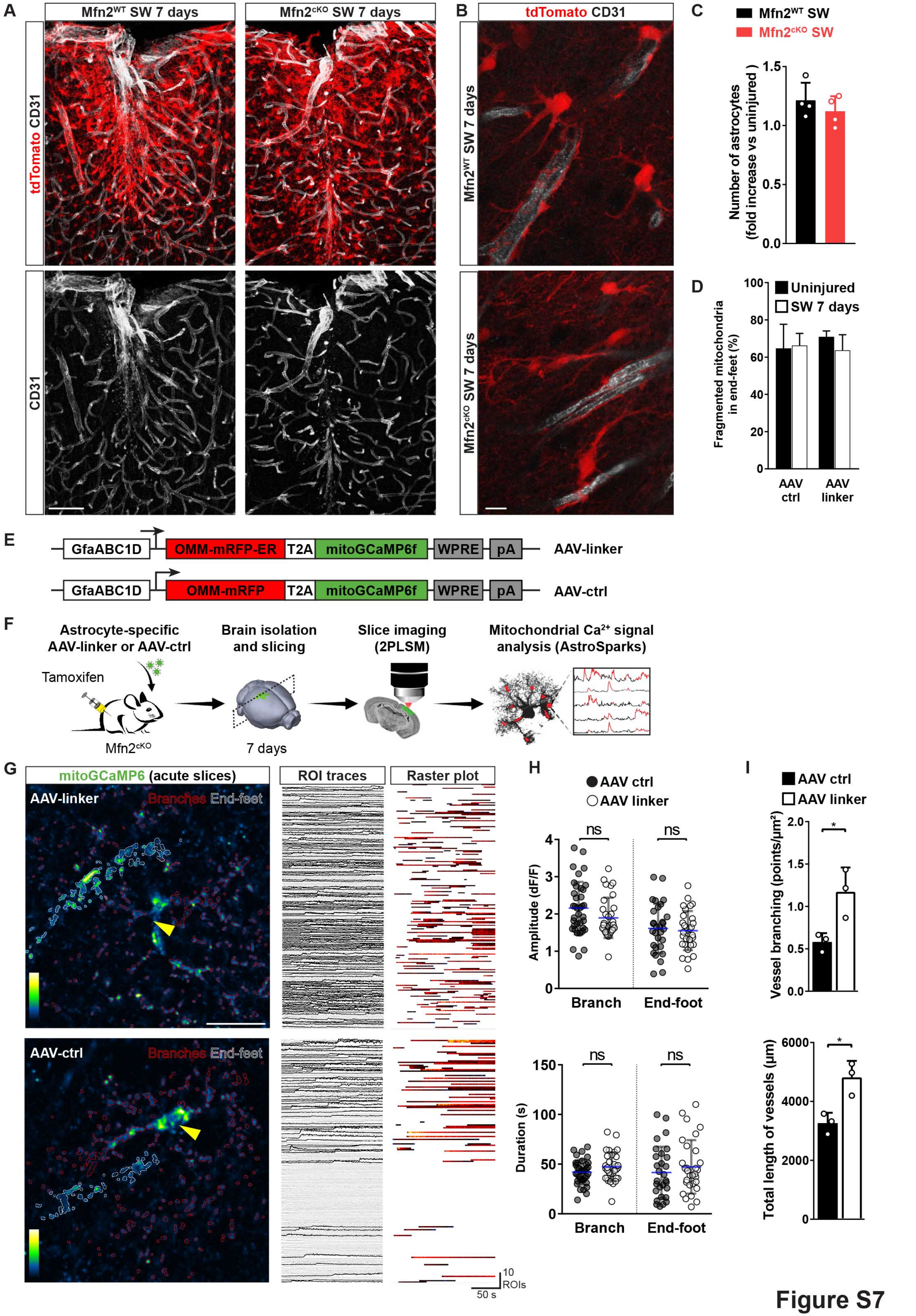
